# Tracking tree demography and forest dynamics at scale using remote sensing

**DOI:** 10.1101/2024.06.11.598435

**Authors:** Robin Battison, Suzanne M. Prober, Katherine Zdunic, Toby D. Jackson, Fabian Jörg Fischer, Tommaso Jucker

**Affiliations:** School of Biological Sciences, University of Bristol, Bristol, BS8 1TQ, UK; CSIRO Environment, Canberra, ACT 2601, Australia; Biodiversity and Conservation Science, Department of Biodiversity, Conservation and Attractions, Kensington, WA 6151, Australia

**Keywords:** competition, growth, LiDAR, mortality, recruitment, remote sensing, topography, tree crown delineation, wildfires

## Abstract

- Capturing how tree growth and survival vary through space and time is critical to understanding the structure and dynamics of tree-dominated ecosystems. However, characterising demographic processes at scale is inherently challenging, as trees are slow-growing, long-lived, and cover vast expanses of land.
- We used repeat airborne laser scanning data acquired over 25 km^2^ of semi-arid, old-growth temperate woodland in Western Australia to track the height growth, crown expansion and mortality of 42,810 individual trees over nine years.
- We found that demographic rates are constrained by a combination of tree size, competition and topography. After initially investing in height growth, trees progressively shifted to crown expansion as they grew larger, while mortality risk decreased considerably with size. Across the landscape, both tree growth and survival increased with topographic wetness, resulting in vegetation patterns that are strongly spatially structured. Moreover, biomass gains from woody growth generally outpaced losses from mortality, suggesting these old-growth woodlands remain a net carbon sink in the absence of wildfires.
- Our study sheds new light on the processes that shape the dynamics and spatial structure of semi-arid woody ecosystems and provides a roadmap for using emerging remote sensing technologies to track tree demography at scale.

## INTRODUCTION

Forest ecosystems face growing pressure on multiple fronts, from increasingly frequent and severe droughts and heatwaves, larger and more intense wildfires and storms, novel pests and pathogens, and human-driven degradation (Senf *et al*., 2018; Canadell *et al*., 2021; Hammond *et al*., 2022; Turner & Seidl, 2023). Understanding how trees are responding to these novel disturbance regimes is critical if we are to forecast how forest dynamics will change over the coming century, and what implications this will have for biodiversity and carbon storage in these ecosystems (McDowell *et al*., 2020; Turner & Seidl, 2023). To achieve this, we need demographic information that allow us to infer and model changes in population dynamics at scale – data that capture how rates of tree growth, mortality and recruitment vary across both space and time (Coomes *et al*., 2014; Fisher *et al*., 2018; Kunstler *et al*., 2021; Needham *et al*., 2022b; Zuidema & van der Sleen, 2022). Ecologists have traditionally relied on networks of permanent field plots to estimate these demographic rates (Lines *et al*., 2010; Ruiz-Benito *et al*., 2013; Kunstler *et al*., 2021; Needham *et al*., 2022b; Piponiot *et al*., 2022). However, while plot networks remain the gold standard to characterise community-level dynamics, they have some inherent limitations when it comes to capturing variation in demographic rates across landscapes. Field surveys are incredibly labour intensive, meaning that most forest plots are small (0.1–1 ha) and cumulatively only cover a tiny fraction of the total forest area (<0.01% even in best case scenarios; Yu *et al*., 2022; Holcomb *et al*., 2023). This makes it challenging to understand how demographic rates vary across environmentally heterogeneous landscapes and in response to large, infrequent disturbances.

Remote sensing offers an intuitive solution to this challenge of tracking large numbers of trees across broad spatial scales (Stovall *et al*., 2019; Brandt *et al*., 2020; Ma *et al*., 2023). In particular, technologies such as airborne laser scanning (ALS, or LiDAR) can be used to build highly accurate and detailed 3D models of both the forest canopy and the underlying terrain (≤1-m resolution) that span thousands of hectares (Jucker, 2022; Lines *et al*., 2022). Unsurprisingly, ALS has become an integral tool for large-area mapping of forest structure and biomass and there is now a growing interest in using repeat ALS acquisitions to quantify forest dynamics at scale (Asner & Mascaro, 2014; Dalponte *et al*., 2019; Dalagnol *et al*., 2021; Nunes *et al*., 2021; Cushman *et al*., 2021; Jucker *et al*., 2023). To date, almost all this work has focused on characterising dynamic processes occurring at a canopy level, such as those associated with the formation, expansion and closure of gaps (Wedeux *et al*., 2020; Dalagnol *et al*., 2021; Nunes *et al*., 2021; Cushman *et al*., 2021; Choi *et al*., 2023). However, in parallel there has also been a push towards developing computational tools to detect individual tree crowns using ALS (Dalponte & Coomes, 2016; Ferraz *et al*., 2016; Cao *et al*., 2023). Benchmarking efforts have shown that algorithms can accurately segment and measure the crown dimensions of canopy-dominant trees (Wang *et al*., 2016; Cao *et al*., 2023), particularly when applied to high-resolution ALS data acquired in open canopy forests. As repeat ALS data become increasingly available, individual-based methods provide a unique opportunity to characterise how rates of tree growth and mortality vary across size-structured populations (Piponiot *et al*., 2022; Brandt *et al*., 2024) and whole landscapes (Duncanson & Dubayah, 2018; Stovall *et al*., 2019; Beese *et al*., 2022; Ma *et al*., 2023).

The appeal of individual-based methods applied to large-scale remote sensing datasets is not just in their ability to inventory huge numbers of trees. By capturing the 3D architecture of individual tree crowns, ALS data also provide a way to characterise tree growth along multiple axes, including height growth and lateral crown expansion (Lines *et al*., 2022). Vertical and horizontal crown growth are rarely recorded in field data, as they are much more complex and time consuming to measure compared to stem diameters. However, they are arguably much more ecologically meaningful when it comes to capturing whole-plant growth strategies and how these vary with tree size, such as hypothesised shifts in biomass allocation away from height growth and towards crown expansion as trees approach maturity (Antin *et al*., 2016; Marziliano *et al*., 2019; Jucker *et al*., 2022; Laurans *et al*., 2024). Similarly, they are much more informative when it comes to understanding competitive interactions for light and space among neighbouring trees (Jucker *et al*., 2015; Taubert *et al*., 2015), as well as providing a way to quantify crown damage and dieback, which are strong predictors of tree mortality (Needham *et al*., 2022a; Zuleta *et al*., 2022). Another appeal of ALS data is that they contextualise the biotic and abiotic environment within which trees grow, such as their local competitive neighbourhood and topographic position within the landscape (Colgan *et al*., 2012; Swetnam *et al*., 2017; Beese *et al*., 2022; Ma *et al*., 2023). This provides an opportunity to not only quantify how demographic rates vary across the landscape, but also ascribe this variation to underlying ecological drivers. Finally, a key selling point of individual-based methods is that they are directly comparable to how we monitor forests on the ground and how we represent them in forest dynamics models. This provides an opportunity to bridge the gap between field and remote sensing forest monitoring programmes and can help reduce major sources of uncertainty in forest dynamics models, such as those associated with tree mortality (Hubau *et al*., 2020; McDowell *et al*., 2020; Pugh *et al*., 2020).

Here we use data from two ALS surveys acquired nine years apart in Australia’s Great Western Woodlands (GWW) to capture the height growth, crown expansion, crown dieback and mortality of individual trees across 2500 ha of old-growth woodland habitat. These semi-arid woodlands are an ideal testbed for using repeat ALS data to quantify tree demographic rates at scale, as they are dominated by a small number of eucalypt species that form sparsely populated stands of single-stemmed trees. By developing a new pipeline for segmenting and matching tree crowns across ALS surveys, we were able to confidently identify and track the dynamics of 42,810 canopy-dominant trees across this landscape. Using these data, we set out to:

1. Determine how growth and mortality rates vary with tree size across the whole population. This allowed us to quantify which cohorts contribute most to biomass gains and losses, as well as characterise how trees adjust their crown growth strategies as they increase in size.
2. Model how tree growth and mortality rates vary across the landscape in relation to fine-scale topography and local neighbourhood competitive environment, so that we may better understand how demographic processes give rise to vegetation spatial patterns in dry forests.
3. Scale up tree-level demographic rates to community-level dynamics in aboveground biomass and canopy 3D structure. In doing so we explored whether temporal changes in canopy 3D structure are predominantly driven by tree growth or mortality, and aimed to determine if these old-growth woodlands currently operate as a net carbon sink or source in the absence of large disturbances from wildfires.

## MATERIALS AND METHODS

### Study system

The study was conducted at the GWW Terrestrial Ecosystem Research Network (TERN) SuperSite (30°12’2’’S, 120°38’55’’E; see Fig. S1 in Supporting Information), one of 16 intensively monitored TERN SuperSites distributed across Australia (Prober *et al*., 2023). Formerly a pastoral lease, since 2007 the land has been managed as a conservation area by the Western Australia Department of Biodiversity, Conservation and Attractions. The site is located on relatively flat terrain (430–475 m.a.s.l.) with a mean annual temperature is 19 °C and the mean annual rainfall is 260 mm. Soils are primarily deep red calcareous loams and clays over a deeply weathered regolith, with a hypersaline, acidic water table at >20 m.

The landscape consists of a mosaic of different vegetation types, with the vast majority covered by old-growth temperate eucalypt woodlands and the rest consisting of mulga scrub (*Acacia* sp.) and heathland (which collectively cover <0.1% of the study area and were masked out of all analyses presented here; see Methods S1 for details). The woodland habitats, which are the focus of this study, are dominated by a small number of obligate-seeder eucalypt species. The most abundant of these is *Eucalyptus salmonophloia*, but *E. salubris, E. transcontinentalis*, and *E. clelandiorum* are also found in relation to subtle variation in soils and topography. Wildfires are the major agent of disturbance in the region (Jucker *et al*., 2023) and are almost always stand replacing, as obligate-seeder eucalypts are highly susceptible to fire and have almost no ability to resprout (Gosper *et al*., 2018). The site has not been affected by wildfires for several hundred years and the woodlands are currently in an old-growth successional stage. This is characterised by open-canopy stands dominated by sparse, large, single-stemmed trees, where localised stand dynamics are initiated by infrequent windthrows and floods which are followed by subsequent recruitment in gaps (Gosper *et al*., 2018). Based on field data collected across three 1-ha forest plots dominated by *E. salmonophloia*, but *E. salubris and E. transcontinentalis*, typical ranges of tree density, basal area and aboveground biomass for old-growth woodlands at this site are 15–34 trees ha^−1^, 4.9–5.1 m^2^ ha^−1^ and 37.9–45.3 Mg ha^−1^, respectively (field data available at: https://field.jrsrp.com).

### Airborne data acquisition and processing

ALS data were acquired at two points in time over a 5×5 km square area centred on the GWW SuperSite (Fig. 1), first in May 2012 and then again exactly 9 years later in May 2021. The 2012 data were collected by Airborne Research Australia (https://www.airborneresearch.org.au) using a motorised glider (Diamond Aircraft, HK36 TTC-ECO) mounted with a RIEGL LMS-Q560 scanner. Flights were conducted at 300 m above ground with a scan angle of ±24°, resulting in a footprint size of 15.0 cm and an average pulse density of 21.4 pulses m^−2^. The 2021 data were acquired by Aerometrex (https://aerometrex.com.au) using a Cessna 404 Titan mounted with a RIEGL VQ-780ii scanner. Flights were conducted at 1100 m above ground with a scan angle of ±30°, resulting in a footprint size of 19.8 cm and an average pulse density of 23.6 pulses m^−2^. During the second ALS survey, high-resolution RGB imagery (20 cm ground sample distance) were also acquired over the study area and provided as an orthomosaic. Georeferenced point clouds for both ALS surveys were provided in LAS 1.2 format and all subsequent processing was performed using a combination of *CloudCompare* (https://www.danielgm.net/cc/), QGIS (https://qgis.org) and R (R Core Team, 2024) using the *lidR* package (Roussel *et al*., 2020).

**Fig. 1:**
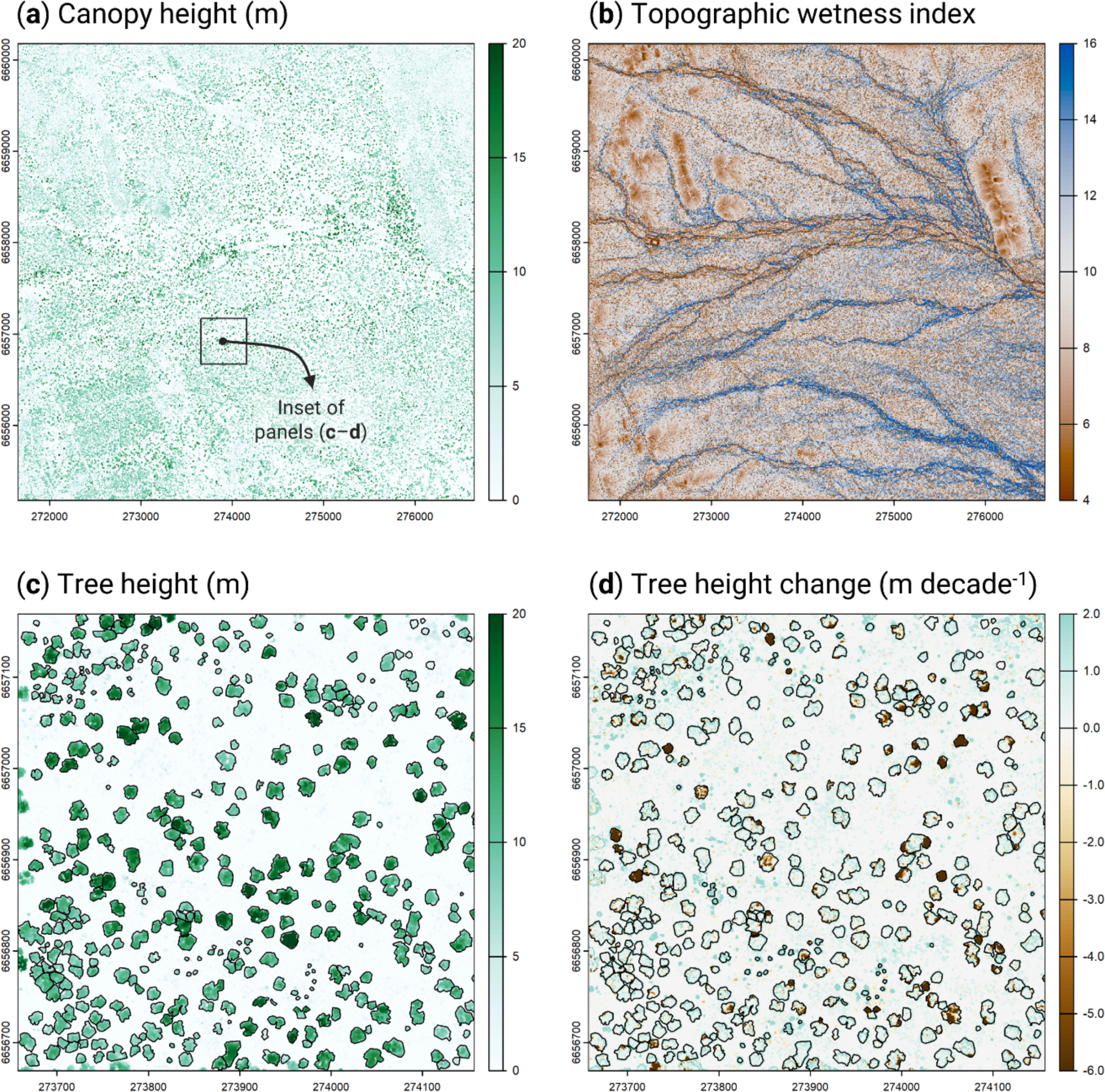
Tracking variation in tree demography across the Great Western Woodland TERN SuperSite using airborne laser scanning (ALS). Panel (**a**) shows the canopy height model (CHM) of the entire study area (5×5 km) generated using the 2012 ALS data, while (**b**) shows variation in topographic wetness index (TWI) across the same extent, from which the major drainage channels of the site can be clearly seen in blue. Panel (**c**) corresponds to a 500×500 m section of the whole CHM (see black inset box in **a**), within which the crowns of individual canopy trees can be clearly seen. Changes in tree height between the two ALS surveys (2012 and 2021) are depicted in (**d**), where trees that died or underwent crown dieback are clearly visible in dark brown.

First, a statistical outlier removal filter implemented in *CloudCompare* was used to identify and remove outlier and duplicate points, following which we standardised the two acquisitions by removing any points with scan angles exceeding ±24°. Each point cloud was then classified into ground and non-ground returns, allowing a normalised point cloud to be produced for both timesteps. From these, a 0.5 m resolution canopy height model (CHM) was then created for both 2012 and 2021 using the pit-free algorithm implemented in the *lidR* package (Khosravipour *et al*., 2014). Alignment between the two CHMs was visually assessed and adjusted in QGIS, at which point we clipped the 2021 CHM to the extent of the 2012 CHM to only retain areas of overlap between the two flights (2500 ha in total). Finally, the 2021 classified point cloud was used to create a 5-m resolution digital terrain model (DTM) of the study area using the triangular irregular network algorithm implemented in *lidR*. All subsequent data processing and analyses were carried out in R using a combination of the *terra (Hijmans et al., 2023)*, *sf* (Pebesma, 2018) and *dynatopmodel* packages (Metcalfe *et al*., 2015).

### Automated tree crown detection and segmentation

Several algorithms have been developed in recent years to automatically detect and delineate individual tree crowns in ALS-derived CHMs (Eysn *et al*., 2015; Dalponte & Coomes, 2016; Aubry-Kientz *et al*., 2019; Cao *et al*., 2023). To determine which of these was most suitable for our study system, we began by manually delineating the crowns of all trees taller than 4 m and with a crown area greater than 9 m^2^ within a 1000×500 m (50 ha) section of the 2012 CHM (*n* = 797 manually delineated crowns in total). Using this as a benchmark, we first compared four alternative crown delineation algorithms implemented in the *lidR* package (*dalponte2016*, *li2012*, *silva2016* and *watershed*) based on several complementary criteria, including: (i) total number of crowns detected, (ii) mean crown area of correctly matched crowns, (iii) percentage of correctly segmented, over-segmented and omitted crowns, (iv) Intersection over Union (IoU, %) of manually delineated and algorithm-segmented crown polygons, and (v) and an F_1_ score obtained by first classifying crowns as either correctly segmented or not using IoU > 50% as a cut-off and then calculating precision and recall as described in (Cao *et al*., 2023). F_1_ scores vary between 0–1, with a value of 0.7 generally being considered good.

Consistent with previous studies (Aubry-Kientz et al., 2019; Eysn et al., 2015), we found that *dalponte2016* performed very well both in absolute terms and relative to the other algorithms we tested (78% correctly segmented crowns; mean IoU = 75%; F_1_ score = 0.92; see Table S1 and Fig. S3 for details). However, despite its robust overall performance, the default implementation of *dalponte2016* exhibited a relatively high degree of over-segmentation (13%), especially for large trees (35% for trees with a crown area >200 m^2^; Fig. S3). To avoid this having an undue effect on our results, we developed a new implementation of the *dalponte2016* routine specifically designed to address this issue of over-segmentation of large trees. This approach is described in detail in Supporting Information (Methods S2), but briefly it involves running the crown segmentation in two stages: first using a broad search window to accurately segment the crowns of large trees (Cao *et al*., 2023), followed by a second pass with a smaller search window to detect any crowns missed in the initial step. Both search windows were defined based on allometric constraints between crown width and height, which we modelled using data from the 797 manually segmented trees (Fig. S2). This two-stage approach considerably improved the performance of the algorithm, increasing overall accuracy to 82% and reducing over-segmentation to 5% (Table S1 and Fig. S4). This new implementation of the *dalponte2016* algorithm was applied to the entire 2012 and 2021 CHMs to segment all trees taller than 4 m and with a crown area greater than 9 m^2^ within the study area.

### Estimating rates of tree height growth, crown expansion, mortality and biomass change

#### Changes in tree height and crown area

To robustly estimate changes in the height and crown area of individual trees, we developed a routine for matching tree crowns detected across both ALS surveys. We began by checking if the crown polygon of a tree delineated in 2012 also contained the centroid of a crown detected in 2021, keeping only those that did. A crown that was only delineated in 2012 would indicate either a tree that died or underwent substantial crown dieback between surveys (or possibly one that was mistakenly omitted by the segmentation routine, but these are a minority; Table S1). Then, to retain only high-quality matches, we removed any instances in which a crown delineated in one of the two ALS surveys contained two or more crown centroids in the other time period. This step was designed to detect instances in which the delineation algorithm mistakenly over-segmented a tree into multiple crowns. While our analysis of the training data suggests over-segmentation errors were infrequent (5%; Table S1), retaining them would bias our estimates of tree growth by introducing extreme values of height and crown area change.

This left us with a subset of crowns for which we have a high degree of confidence that the same tree was detected at both time points. From these we further removed any trees that underwent a ≥30% decrease in height and/or crown area, which were classified as dead or having undergone severe dieback (Duncanson & Dubayah, 2018; Ma *et al*., 2023). All remaining matched crowns were used to estimate changes in individual tree height (m decade^−1^) and crown area (m^2^ decade^−1^) between the two ALS acquisitions. Tree height was defined as the maximum value of the CHM within the crown polygon, which is analogous to how it is typically measured in field surveys (Jucker *et al*., 2022); although we note that taking the mean value of the CHM yielded very similar results; Fig. S5).

#### Tree mortality and crown dieback

Two complementary approaches were used to identify trees that died or underwent pronounced crown dieback (hereafter treated as a single, combined category) between the two ALS surveys (Fig. 1d). First, we used the set of matched crowns described above to screen for trees that decreased in height and/or crown area by ≥30% throughout the study period and classified these as dead or having experienced severe dieback (Duncanson & Dubayah, 2018; Ma *et al*., 2023). Second, to capture trees that completely disappeared between the two surveys (i.e., those identified only in the 2012 scan), we overlayed all unmatched crowns from 2012 onto both CHMs and measured their change in height from one time point to the next. Applying the same threshold as before, all trees that decreased in height by ≥30% were classified as dead.

Note that the inverse approach could be used to estimate rates of tree recruitment into the population (i.e., trees exceeding a height ≥4 m and crown area ≥9 m^2^ only in 2021). However, we chose not to implement this into our analysis, as trees in these smaller size classes exhibited the highest rates of both height growth and mortality (see Results). As a result, over the course of the 9-year period separating the two ALS scans, trees could have surpassed the minimum size threshold only to then die before the second survey took place, causing true recruitment rates to be substantially underestimated.

#### Aboveground biomass stocks and changes

To estimate the aboveground biomass (*AGB*, in kg) of individual trees detected in the 2012 and 2021 ALS surveys, we followed a two-stage approach based on the framework developed in Jucker *et al*. (2017). First, we estimated each tree’s diameter at breast height (*DBH*, in cm) based on its height (*H*, in m) and crown diameter (*CD*, in m) using the following allometric model: *DBH* = 0.519 × (*H* × *CD*)^0.890^ × *CH*_*DBH*_, where *CF_DBH_* = 1.002 and corresponds to the Baskerville correction factor. *H* and *CD* were derived from the segmented tree crown polygons, with *CD* calculated from crown area assuming a circular crown. This allometric model was developed using data from angiosperm trees growing in woodland and savanna ecosystems of Australasia (Jucker *et al*., 2017).

We then estimated each tree’s aboveground biomass using an allometric model developed specifically for eucalypt trees in Australia (Paul *et al*., 2016): *AGB* = 0.133 × (*DBH*)^2.375^ × *CF_AGB_*, where *CF_AGB_* = 1.067. Each tree’s aboveground biomass change was then calculated as the difference in *AGB* between the two ALS surveys (expressed in kg decade^−1^). For these calculations, trees that were classified as having died or undergone severe crown dieback were assumed to have lost all their biomass (*AGB_2021_* = 0).

### Characterising size-dependent rates of tree growth, mortality and biomass change

To build an overall picture of population-level trends in tree demographic rates across the landscape and how they vary with tree size, we aggregated the tree-level data on height growth, crown expansion and mortality according to size class. As a measure of initial tree size, for each tree we used the data from 2012 to calculate the product of its height and crown diameter (*H*×*CD*, in m^2^), which previous work has shown to be strongly related to a tree’s total aboveground biomass (Jucker *et al*., 2017; Ma *et al*., 2023). Trees were then assigned to one of 10 size classes by binning values of *H*×*CD* into logarithmic bins of equal width (logarithmic binning was chosen to better capture the right-skewed distribution of the data). Each size class was represented by >1750 trees (range = 1761–7382 trees per size class). To test how rates of tree height growth and crown expansion vary with tree size, we fit ANOVAs to compare mean values of both growth metrics across the 10 size classes. Similarly, to characterise how tree mortality rates change with tree size, we used logistic regression (generalized linear model with binomial errors and logit link) to estimate the probability of a tree being alive (0) or having died/experienced severe dieback (1) by the time of the second ALS survey for each of the 10 size classes.

To understand how variation in growth rates and mortality translate to changes in aboveground biomass, for each size class we calculated the total biomass gains of trees that survived, the biomass losses of those that died, as well as the net biomass change between the two ALS surveys (each expressed in Mg ha^−1^ decade^−1^). For this analysis, trees were binned into 10 percentile size classes (rather than equal width bins), so that each size class contained the same number of trees (*n* = 4281). This allowed us to directly compare the contribution of each size class to the aboveground biomass dynamics of the whole population.

### Drivers of landscape-scale variation in tree demographic rates

To determine what processes shape variation in tree demographic rates across the landscape, we modelled variation in height growth, crown expansion, and mortality and dieback as a function of several intrinsic and extrinsic factors. First, to capture how growth allocation to height, crown expansion and risk of mortality vary in relation to tree size and life stage (Ruiz-Benito *et al*., 2013; Jucker *et al*., 2022), we included initial tree size (i.e., *H*×*CD* in 2012) as a model predictor. Next, to account for the effects of neighbourhood competition for light, water and nutrients on tree growth and mortality (Canham *et al*., 2004; Ma *et al*., 2023), we used the 2012 CHM to calculate a metric of local competitive environment for each tree. Specifically, a 25-m buffer was added around the perimeter of each delineated crown, within which we calculated the mean height of the CHM as a proxy of neighbourhood competition (Ma *et al*., 2023). Then, to test how demographic rates vary in relation to local topography (Jucker *et al*., 2018), we used the *dynatopmodel* package to calculate topographic wetness index (TWI) from the DTM (Fig. 1b) and then assigned a value of TWI to each tree based on their location. TWI describes how water flows and accumulates across the landscape, thus providing an indicator of local soil water and nutrient availability for plants (Kopecký *et al*., 2021). Finally, to control for differences in pulse density between ALS surveys which could otherwise bias estimates of growth and mortality (Roussel *et al*., 2017), we included the difference in pulse density between the two scans as a model predictor (Fig. S6). This accounts for the fact that we would overestimate growth rates of trees scanned with a higher pulse density in 2021 simply by virtue of the more intense sampling of their crowns (and *vice versa* for 2012).

Multiple linear regressions were used to model variation in individual tree height growth and crown expansion rates, while probability of mortality was modelled using logistic regression (generalized linear model with binomial errors and logit link). Initial tree size (*H*×*CD*) was log-transformed to account for its strongly right-skewed distribution. Additionally, to allow direct comparisons between model coefficients, all predictors were scaled and centred to have a mean of 0 and a standard deviation of 1 prior to model fitting.

### Mapping spatial variation in demographic rates and its impacts on biomass dynamics and canopy 3D structure

To characterise spatial variation in demographic rates across the landscape, we mapped mean rates of tree height growth, crown expansion and tree mortality (% of stems that died or underwent severe crown dieback) across the GWW SuperSite. Individual tree-level data were aggregated and mapped at a 1-ha resolution across the 5×5 km study area (*n* = 2491 ha, after masking out 9 ha dominated by mulga habitat). This allowed us to visualise spatial patterns in demographic rates across the landscape and quantify the degree to which growth and mortality rates co-vary spatially. Then, to better understand how tree growth and mortality combine to drive aboveground biomass dynamics, we used the same approach to also map net changes in aboveground biomass across the landscape. In doing so we explored whether biomass dynamics are more strongly constrained by tree growth or mortality, and assessed if these old-growth woodlands are currently operating as a net carbon sink or source.

To complement these individual tree-based estimates of biomass dynamics, we also calculated two independent measures of changes in canopy 3D structure, which we derived directly from the CHMs at 1-ha resolution. The first was a measure of change in canopy cover at 4-m aboveground between 2021 and 2012 (Δ_cover_, expressed as % decade^−1^). A 4-m height threshold was chosen to match that used to include trees in our segmentation routine. The second was an estimate of change in canopy volume (Δ_vol_, expressed as m^3^ decade^−1^) over the same period, where canopy volume was calculated by multiplying the height of each CHM pixel by its area and then summing across the 1-ha grid. By quantifying the degree of spatial correlation between aboveground biomass change and Δ_cover_ and Δ_vol_, we were able to determine how well changes in canopy 3D structure can be inferred when the underlying demographic rates are known.

## RESULTS

Across the GWW SuperSite, we identified a total of 55,043 tree crowns in 2012 and 56,833 in 2021 (22.1 and 22.8 trees ha^−1^, respectively). Of these, we were able to confidently match 42,810 tree crowns across the two surveys, 3524 of which were classified as having died or experienced severe dieback between the two ALS surveys (10.0% decade^−1^). In 2012, matched trees had a mean height of 10.9 m (interquartile range = 8.1–13.1 m; 99^th^ percentile = 20.9 m) and a mean crown area of 122.8 m^2^ (interquartile range = 46.8–171.8 m^2^; 99^th^ percentile = 436.3 m^2^). By 2021, trees that survived had grown an average of 0.20 m decade^−1^ in height (interquartile range = 0.07–0.37 m decade^−1^) and expanded their crowns by 13.6 m^2^ (interquartile range = 0.3–24.7 m^2^ decade^−1^). Estimated height growth rates were noticeably larger than instrument measurement errors, which we estimated by comparing the 2012 and 2021 DTMs (root mean square error = 0.03 m).

### Size-dependent variation in tree demographic rates and biomass dynamics

Mean rates of tree height growth, crown expansion and mortality all varied considerably with tree size (Fig 2). Tree height growth peaked in small trees (0.50 m decade^−1^) and decreased progressively thereafter to just 0.05 m decade^−1^ in the largest size classes (Fig. 2a). By contrast, crown area showed the opposite pattern (Fig. 2b), with larger trees investing more than twice as much as small trees in crown expansion (16.0 and 7.3 m^2^ decade^−1^, respectively). Together, these opposing growth trends highlight how trees progressively shift their woody biomass allocation away from tree height growth and towards crown expansion as they become larger (Fig. S7). As for risk of mortality and severe crown dieback, this initially decreased sharply with tree size before plateauing to around 7% decade^−1^ in medium and large-sized trees (Fig. 2c). Overall, trees in the smallest size class were around four times as likely to die or undergone dieback (23.7% decade^−1^) as those in the largest size class (6.7% decade^−1^).

**Fig. 2:**
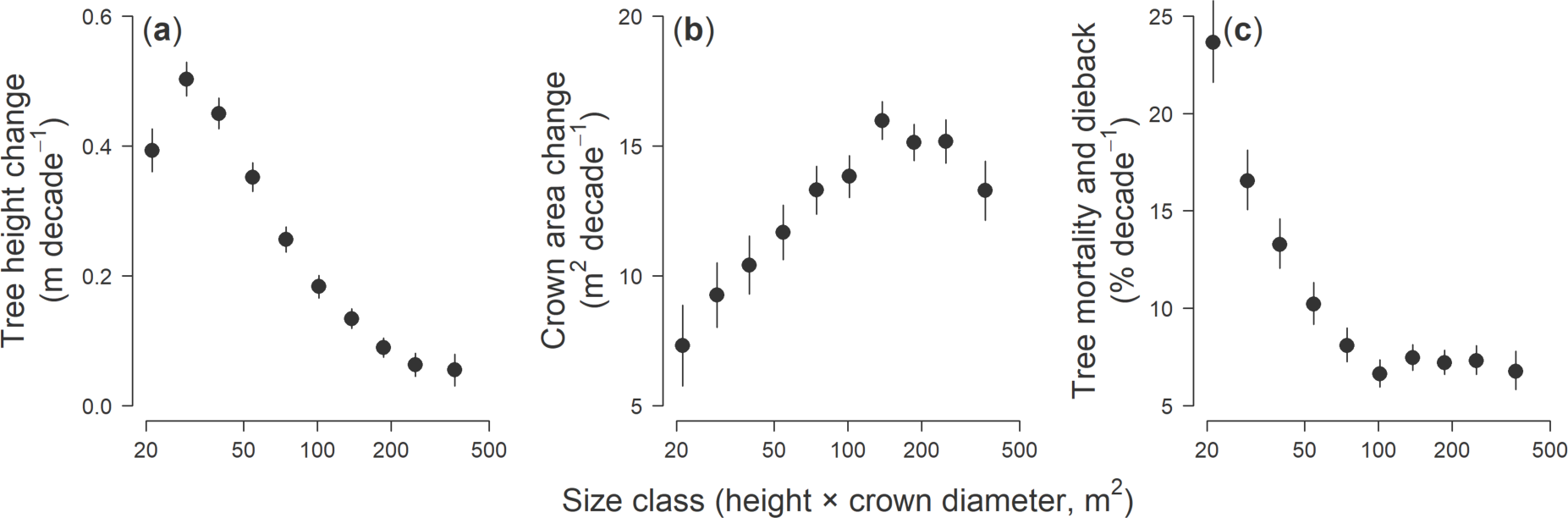
Variation in mean (**a**) tree height growth, (**b**) crown area growth and (**c**) tree mortality and crown dieback rates among tree size classes. Error bars correspond to 95% confidence intervals of the mean for each size class. Tree size was expressed as the product of tree height and crown diameter, which we grouped into 10 equal width bins. Each size class in the analysis is represented by over 1750 trees. Trees were classified as dead or having undergone dieback if they exhibited a decrease in height and/or crown area of ≥30% between the two ALS surveys.

When growth and mortality rates were translated to changes in aboveground biomass, we again observed strong differences among size classes (Fig. 3). In terms of stocks, most of the aboveground biomass is stored in large trees, with the largest 10% of trees accounting for 43.3% of the aboveground biomass, while the smallest 50% only make up 8.5% (Fig. 3a). A similar picture emerged when looking at rates of biomass gains and losses over time, both of which increased continuously with tree size (Fig. 3b). Across the whole population, aboveground biomass gains associated with growth (+2.9 Mg ha^−1^ decade^−1^) were greater than losses driven by mortality and crown dieback (−1.9 Mg ha^−1^ decade^−1^). Consequently, net changes in aboveground biomass were positive (+1.0 Mg ha^−1^ decade^−1^), suggesting that the woodland as a whole is currently operating as a net carbon sink. When broken down by size class we found that it was medium-sized trees that contributed most to this net sink (net aboveground biomass change of trees in the 30–70% size percentile = +0.62 Mg ha^−1^ decade^−1^). By contrast, a combination of decreasing growth rates and progressively greater losses in biomass linked to the mortality of large-stature trees meant that contributions to net biomass gains declined with tree size and even became negative in the largest size class (−0.10 Mg ha^−1^ decade^−1^).

**Fig. 3:**
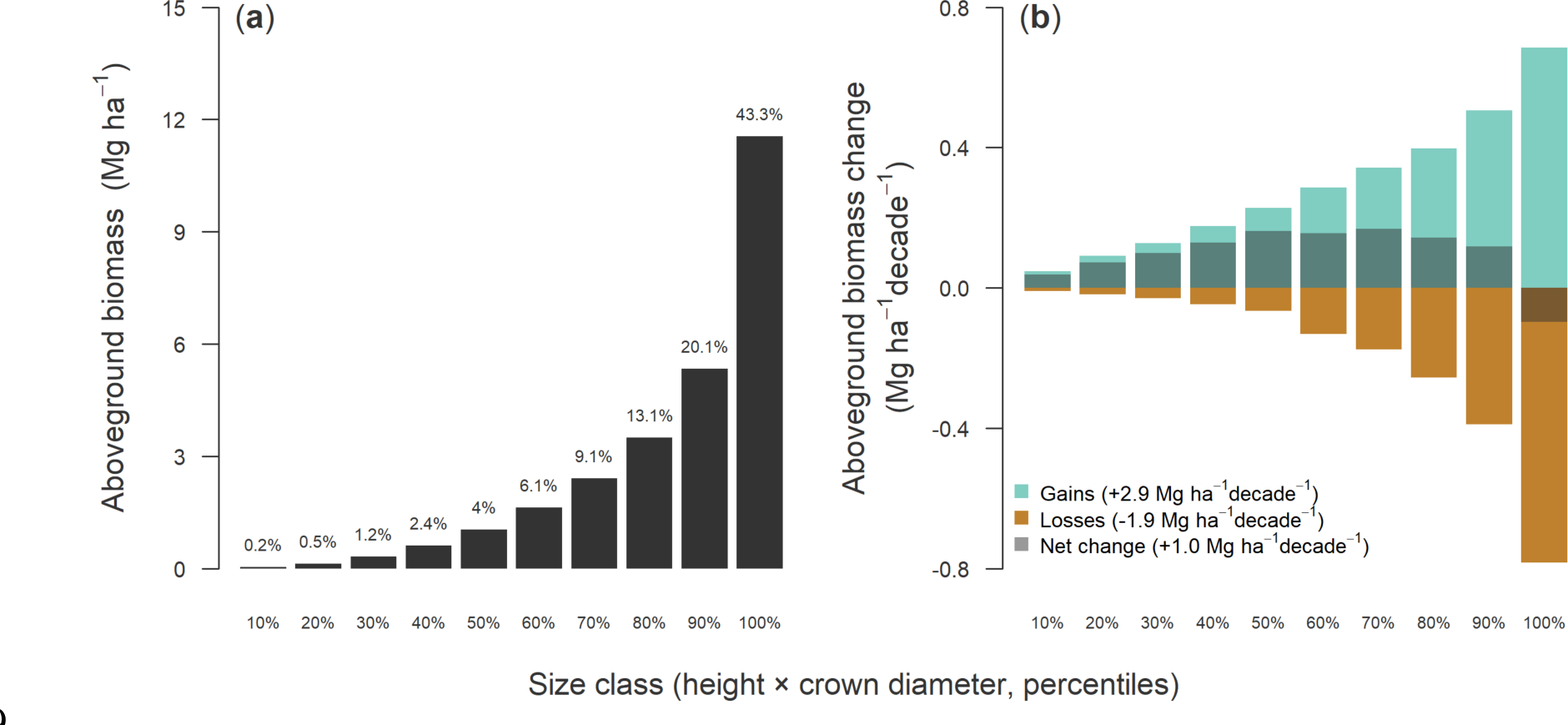
Variation of (**a**) aboveground biomass stocks and (**b**) aboveground biomass dynamics among tree size classes. Trees were binned into 10 percentile size classes based on the product of their height and crown diameter, so that every bin contains the same number of trees. In (**a**) each bar corresponds to the cumulative aboveground biomass of trees belonging to a particular size class at the time of the first survey in 2012. The values reported at the top of each bar denote the proportion of the total aboveground biomass stored in each size class, averaged across the entire study site (mean total aboveground biomass = 26.7 Mg ha^−1^). In (**b**) bars indicate both biomass grains resulting from tree growth (green) and losses associated with mortality and crown dieback (brown) across each size class. The darker shaded area of each bar corresponds to the net change in aboveground biomass (i.e., gains - losses) for each size class, which was positive (dark green) for all expect the largest size class of trees (dark brown). Values reported in the legend of panel (**b**) are estimated total gains (+2.9 Mg ha^−1^ decade^−1^), losses (−1.9 Mg ha^−1^ decade^−1^) and net change in aboveground biomass (+1.0 Mg ha^−1^ decade^−1^) across all size classes combined.

### Drivers of landscape-scale variation in tree demographic rates

Both topography and local competitive environment emerged as important drivers of variation in tree growth and mortality across the landscape (Fig. 4), each exerting an effect that was comparable in magnitude to that of tree size (Fig. 5). In particular, both tree growth and probability of survival increased with TWI. After accounting for the effects of tree size, neighbourhood height and pulse density, predicted height growth was five times higher for trees in areas of the landscape with high TWI (0.26 m decade^−1^ at 95^th^ percentile of TWI) compared to ones with low TWI (0.05 m decade^−1^ at 5^th^ percentile of TWI). Similarly, rate of crown expansion increased from 11.3 to 16.5 m^2^ decade^−1^ and probability of mortality decreased from 7.2 to 5.9% decade^−1^ when transitioning from low to high TWI (brown *vs* blue curves in the lefthand panels of Fig. 5).

**Fig. 4:**
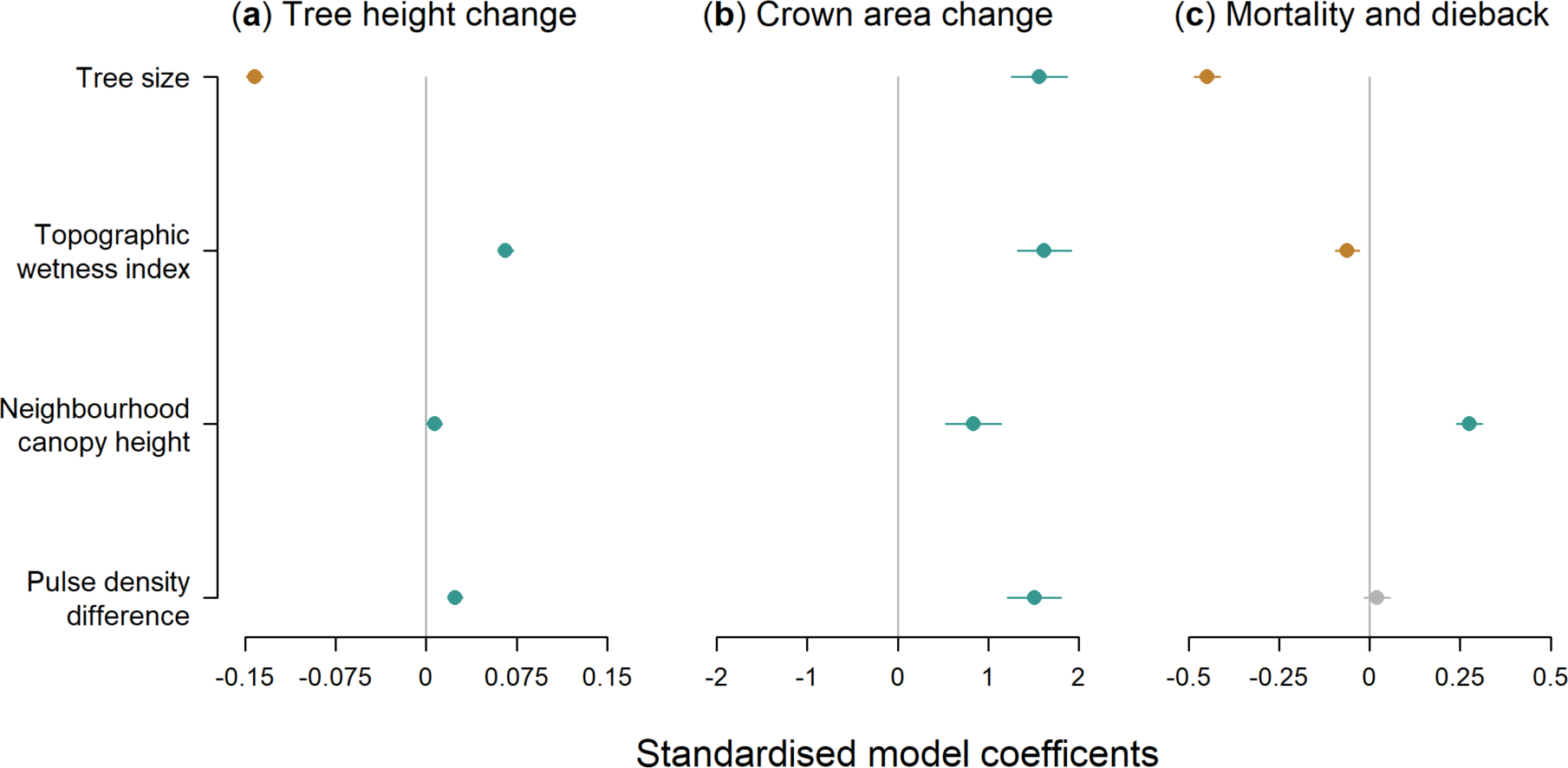
Drivers of variation in (**a**) tree height change (**b**) crown area change and (**c**) risk of mortality and crown dieback across the Great Western Woodland SuperSite between 2012 and 2021. Points are standardized model coefficients, with error bars corresponding to 95% confidence intervals. Significantly positive and negative coefficients are shown in green and brown, respectively, while those for which the 95% confidence intervals overlap with zero are shown in grey. Tree size was defined as the product of tree height and crown diameter, whole pulse density difference corresponds to the change in pulse density between the 2021 and 2012 airborne laser scanning surveys (where positive values indicate trees that were sampled with a greater pulse density in the second survey).

**Fig. 5:**
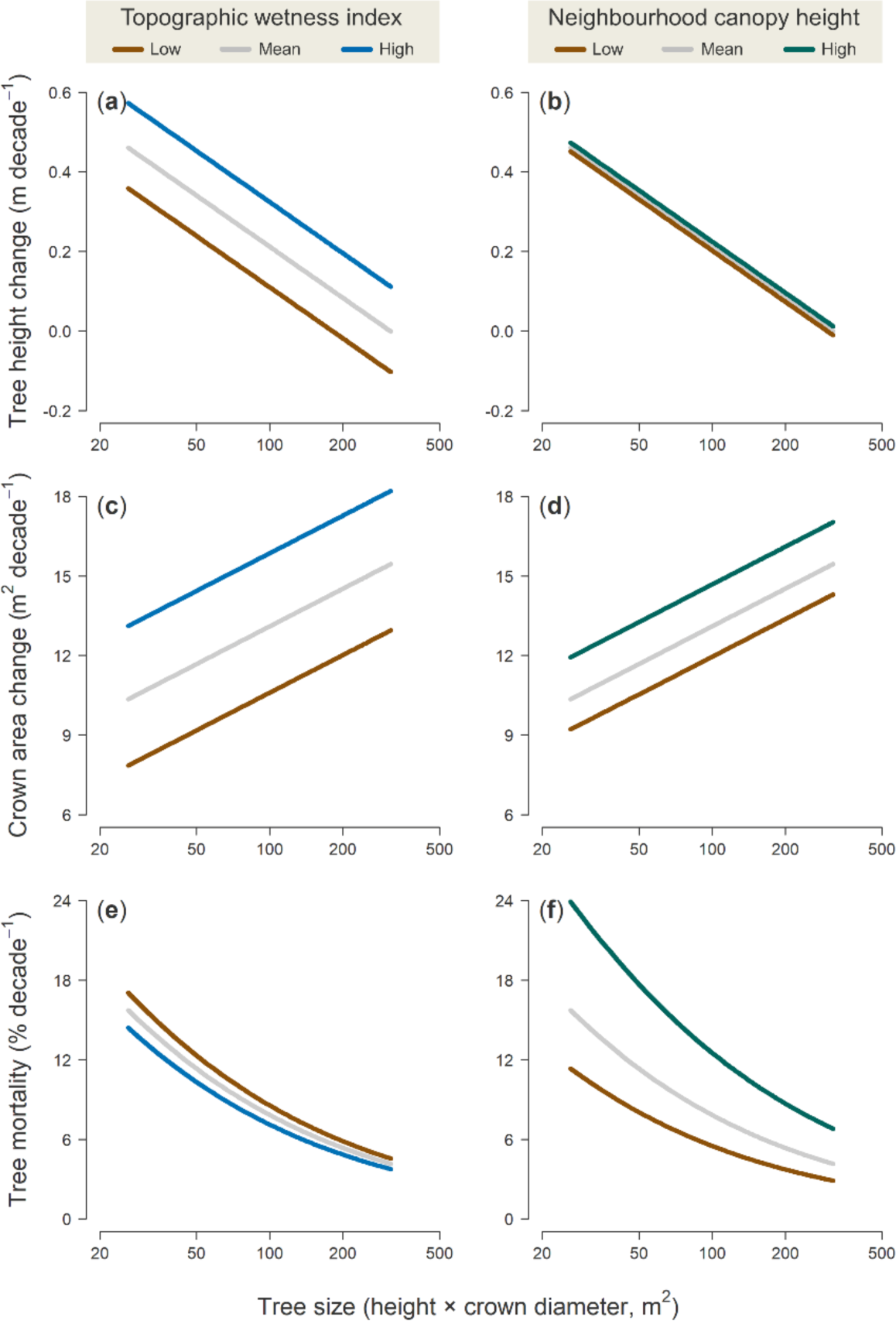
Influence of topographic wetness index (TWI, lefthand panels) and neighbourhood canopy height (righthand panels) on predicted rates of (**a–b**) tree height change (**c–d**) crown area change and (**e–f**) risk of mortality and crown dieback. Curves correspond to predicted values obtained from the multiple regression models shown in Fig. 4, where pulse density difference was set to 0 and tree size was allowed to vary from the 5–95^th^ percentile of the data. For TWI, we predicted values of each demographic rate for trees growing at low (5^th^ percentile, brown curves), mean (grey curves) and high (95^th^ percentile, blue curves) of TWI, while keeping neighbourhood canopy height fixed at its mean value. The same procedure was used to predict variation in demographic rates in relation to neighbourhood canopy height.

Tree growth was also faster in taller stands, especially in the case of crown expansion which for the average surviving tree increased from 12.6 to 15.3 m^2^ decade^−1^ when comparing trees growing in short (5^th^ percentile) *vs* tall (95^th^ percentile) neighbourhoods (brown *vs* green curves in Fig. 5d). However, the opposite was true for mortality, which more than doubled for trees in tall neighbourhoods relative to ones in short, open stands (10.6% vs 4.6% decade^−1^; Fig. 5f). Finally, our analysis also confirmed the importance of statistically controlling for local differences in pulse density among scans, as we found that both height growth and crown expansion were overestimated in trees sampled at higher densities in 2021 (Fig. 4).

### Landscape-scale variation in tree demographic rates and its impacts on biomass dynamics and canopy 3D structure

We observed considerable spatial variation in tree demographic rates aggregated at 1-ha scale (Fig. 6a–c), with tree height change ranging from −0.25–0.65 m ha^−1^ decade^−1^, crown area changes from 0.1–32.8 m^2^ ha^−1^ decade^−1^, and probability of mortality from 0–29.7% ha^−1^ decade^−1^ (values are 5^th^ and 95^th^ percentile of the data shown in Fig. 6a–c). Rates of tree height growth and crown expansion were moderately positively correlated with one another (*ρ* = 0.33). By contrast, we found only a very weak negative relationship between mortality and both height and crown area growth (*ρ* = −0.10 and −0.14, respectively), indicating that these demographic axes were largely decoupled at the landscape scale.

**Fig. 6:**
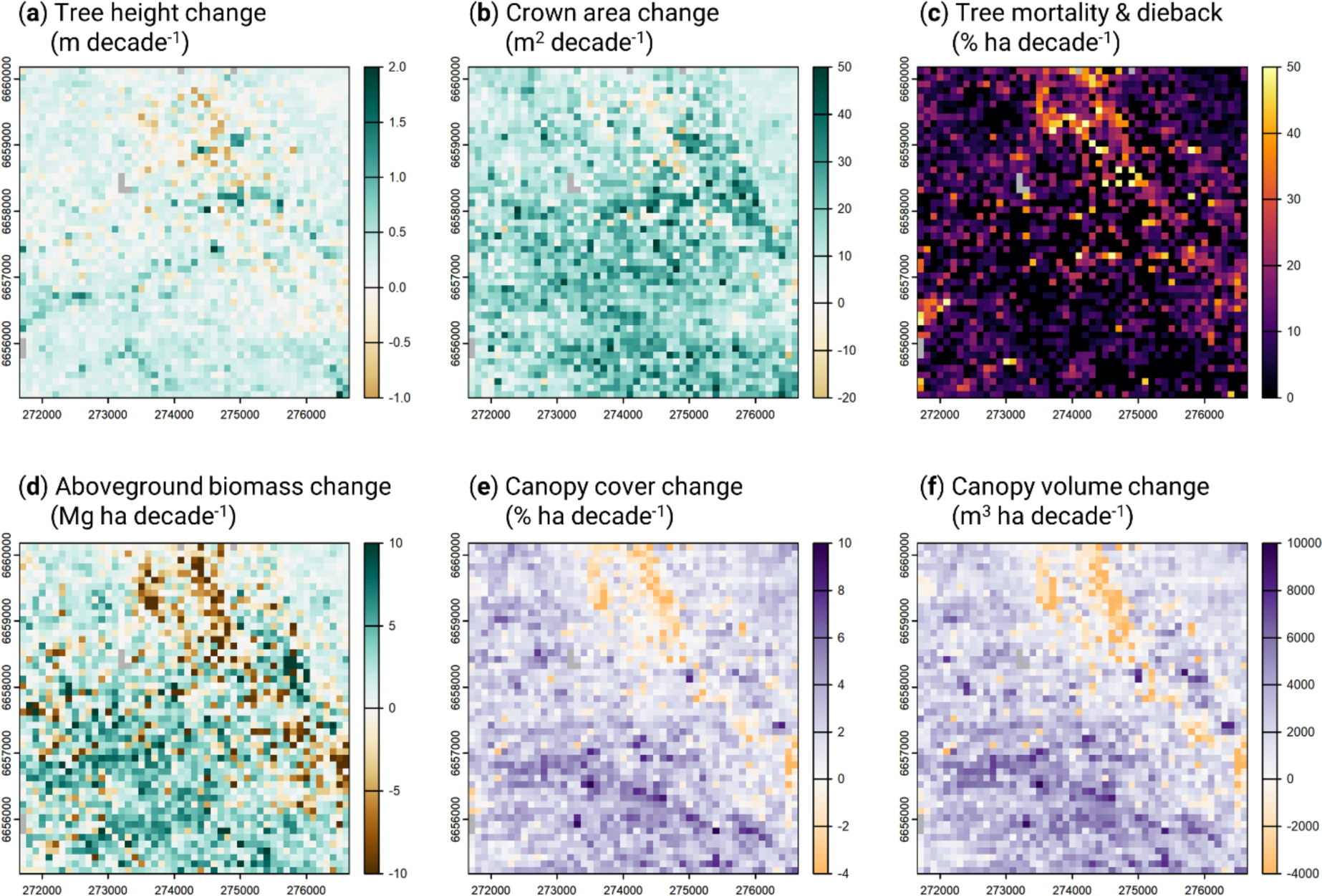
Spatial variation in rates of (**a**) tree height change, (**b**) crown area change, (**c**) tree mortality and dieback, and (**d**) aboveground biomass change across the Great Western Woodland SuperSite between 2012 and 2021. Panels (**a**–**d**) were generated by aggregating the individual tree-level data at 1-ha resolution across the 5×5 km study area (*n* = 2491 ha, after masking out 9 ha dominated by mulga habitat, shown in grey in the maps). For panels (**a**–**b**) only trees that were classified as alive at both census periods were used to generate the maps. For comparison, rates of (**e**) canopy cover change (defined as % ground cover at 4 m aboveground) and (**f**) canopy volume change across the site are also shown. These two canopy-level metrics were calculated directly from the 2012 and 2021 canopy height models, without delineating and tracking changes in individual tree crowns through time.

On average, aboveground biomass stocks increased by 1.0 Mg ha^−1^ decade^−1^ (+3.4%), from 26.7 Mg ha^−1^ in 2012 to 27.6 Mg ha^−1^ in 2021. However, both biomass stocks (6.3–54.8 Mg ha^−1^) and rates of net biomass change over time (−6.9–6.7 Mg ha^−1^ decade^−1^) varied considerably across the landscape (Fig. 6d). Areas that experienced faster rates of tree height growth (*ρ* = 0.35) and crown expansion (*ρ* = 0.47) also had the greatest increases in aboveground biomass, but in general spatial variation in biomass change was most strongly tied to rates of tree mortality (*ρ* = −0.55).

Over the same time period we also found substantial variation in rates of canopy cover (Δ_cover_) and volume change (Δ_vol_) across the landscape (Fig. 6e–f). At the time of the first ALS survey in 2012, mean canopy cover at 4-m aboveground was 24.0% ha^−1^ and mean canopy volume was 25,172 m^3^ ha^−1^. On average, by 2021 both metrics had increased by approximately 7.8% across the landscape, with canopy cover reaching 25.9% ha^−1^ and canopy volume 27,130 m^3^ ha^−1^. But there were also substantial portions of the landscape where both canopy cover and volume decreased relative to 2012. In particular, we observed a very distinct and large diagonal track (approximately 3 km long and 200 m wide) in the northeast corner of the site where most of these losses were concentrated (see orange pixels in Fig. 6e–f). This same pattern is also clearly visible in the tree-level growth, mortality and biomass change maps (Fig. 6a–d), indicating that these two independent approaches to quantifying canopy dynamics are spatially coherent. Specifically, we found that both Δ_cover_ and Δ_vol_ were closely correlated with estimated rates of aboveground biomass change across the landscape (*ρ* = 0.62 and 0.69, respectively).

## DISCUSSION

### Gaining new perspective on tree demography using remote sensing

Using high-resolution repeat ALS data, we were able to quantify rates of tree height growth, crown expansion, dieback and mortality across tens of thousands of trees and map their variation across an entire landscape. This new approach to tracking tree demography from above using remote sensing enabled us to gain unique insights into the process that shape forest dynamics at scale in a way that would have been impossible from the ground.

From a resource allocation perspective, our results showed a clear pattern of trees shifting their growth strategies as they became larger (Fig. 2 and Fig. S7). When small, trees initially invested primarily in height growth, presumably to establish their crowns in the canopy (Laurans *et al*., 2024) and as a strategy to reduce ladder fuels and fire spread (Gosper *et al*., 2018). But as they grew larger, trees began progressively increasing their growth allocation towards crown expansion, with height growth decreasing to almost nothing in the largest size classes. This strategy appears to be common across forest types (Antin *et al*., 2016; Marziliano *et al*., 2019), but makes particularly good sense in open and dry environments such as the GWW. Here being too tall needlessly increases the risk of drought-induced embolism (Olson *et al*., 2018) as there is little to be gained from a light competition perspective. Instead, a more efficient strategy for maximizing light interception is to expand one’s crown outwards. Similar growth patterns have also been observed in African savannas, although here the initial investment in height growth has been attributed not just to escaping fire risk but also herbivory (Moncrieff *et al*., 2011). More generally, by characterizing how growth allocation strategies vary with tree size, we can better understand how static allometric constraints between tree height and crown size emerge as a result of dynamic growth processes (Fischer *et al*., 2019; Jucker *et al*., 2022; Lines *et al*., 2022; Laurans *et al*., 2024). This is a critical step for building more realistic models of forest dynamics that accurately represents the 3D structure of forest canopies (Fischer *et al*., 2020; Jucker, 2022).

From a demography perspective, one the biggest contributions that remote sensing technologies such as repeat ALS can offer is the ability to comprehensively characterize rates and patterns of tree mortality (Duncanson & Dubayah, 2018; Stovall *et al*., 2019; Ma *et al*., 2023). Growing evidence suggests that rates of tree mortality are on the rise globally (Senf *et al*., 2018; McDowell *et al*., 2020; Hartmann *et al*., 2022; Hammond *et al*., 2022) with major consequences for the ability of forests to store carbon (Johnson *et al*., 2016; Pugh *et al*., 2020). However, mortality remains the largest source of uncertainty when it comes to projecting future changes in forest structure and dynamics (Hubau *et al*., 2020; McDowell *et al*., 2020; Pugh *et al*., 2020). This is because tree mortality is an infrequent and stochastic processes that is inherently hard to capture using field inventories. Previous efforts to use remote sensing to address this challenge have mostly focused on canopy-scale disturbances, tracking changes in canopy cover or gap dynamics (Chambers *et al*., 2013; Senf *et al*., 2018; Cushman *et al*., 2021, 2022). A major advantage of our approach is the ability to track mortality and crown dieback at the individual tree level across entire landscapes (Fig. 2c and Fig. 6c). This not only allows direct comparisons with field data, but also provides an intuitive and robust way to scale up from individuals to community-level patterns (Fig. 6).

Across the landscape, average rates of tree mortality were 1.0% yr^−1^, which is consistent with baseline rates documented in other forest types (Lines *et al*., 2010; Ruiz-Benito *et al*., 2013; Stovall *et al*., 2019; Esquivel-Muelbert *et al*., 2020; Hartmann *et al*., 2022). As is typically observed, mortality was highest for small trees and then declined sharply as trees grew larger (Fig. 2c). However, we did not observe a U-shaped pattern of mortality, where survival rates decline in the largest size classes due to greater exposure to disturbances such as wind, lightning and drought (Coomes & Allen, 2007; Lines *et al*., 2010; Coomes *et al*., 2012; Stovall *et al*., 2019). Instead, mortality plateaued at around 0.6–0.7% yr^−1^ in medium and large-sized trees, a pattern that has been reported previously in other Mediterranean woodlands (Ruiz-Benito *et al*., 2013).

When growth and mortality rates were integrated and converted into aboveground biomass gains and losses, differences between size classes became even more apparent (Fig. 3). Similarly to previous work, we found that most of the aboveground biomass is stored in a relatively small number of large trees (Fig. 3a), although this disparity is not as pronounced as in tropical rainforests where the largest 1% of trees can exceed 60–70% of the total standing biomass (Lutz *et al*., 2018). Trees in the largest size classes also contributed most to aboveground biomass gains (green bars in Fig. 3b), as for a big tree even a small increase in height and/or crown size translates into large gains in terms of biomass (Stephenson *et al*., 2014; Jucker *et al*., 2017). However, these gains were partially or even completely offset by progressively greater losses in biomass associated with the mortality and dieback of large trees (brown bars in Fig. 3b), so much so that trees in top 10% for size actually lost more biomass than they gained over the course of the study. Instead, it was mid-sized trees – where rates of crown expansion peaked and mortality was at its lowest (Fig. 2) – that contributed most to net gains in live aboveground biomass across the landscape. This underscores the importance of being able to partition biomass gains and losses across size-structured populations in order to fully understand their dynamics (Piponiot *et al*., 2022; Zuidema & van der Sleen, 2022; Yu *et al*., 2024).

### Linking tree demography to vegetation dynamics and 3D structure

Rates of tree growth and mortality varied noticeably across the landscape in relation to both competitive environment and topography (Fig. 6), the effects of which were similar in magnitude to those of tree size (Figs 4–5). In terms of topography, trees were more likely to have faster rates of vertical and horizontal crown growth, as well as higher likelihood of survival, in area of the landscape that retain more water (high TWI). This is what we would expect in a semi-arid ecosystem such as the GWW, where water is likely the primary limiting factor to tree growth, recruitment and survival (Boisvenue & Running, 2006). It also helps explain previous work showing how vegetation structural attributes such as height, cover and biomass often vary considerably in response to seemingly subtle changes in terrain elevation and slope, typically decreasing from valley bottoms to ridges (Colgan *et al*., 2012; Swetnam *et al*., 2017; Jucker *et al*., 2018; Muscarella *et al*., 2020). The ability of ALS to concurrently map both the 3D structure of the vegetation and the underlying terrain is a major asset of this technology for ecological research (Jucker *et al*., 2018; Muscarella *et al*., 2020). In terms of the effects of local competition, faster rates of tree growth in more densely vegetated areas were balanced out by noticeably greater risk of mortality, particularly for trees in smaller size classes (Fig. 5f). This effect of neighbourhood crowding on tree mortality is consistent with previous work showing that competition for light and water are among the primary drivers of mortality in smaller sized trees (Coomes *et al*., 2012; Ruiz-Benito *et al*., 2013).

More generally, capturing how tree demographic rates are shaped by both topography and local competitive environment allowed us to shed light on the processes that give rise to the clustered distribution of vegetation across the landscape (Fig. 1). These spatial patterns are a hallmark of semi-arid ecosystems (Rodriguez-Iturbe *et al*., 2019; Veldhuis *et al*., 2022) and are increasingly used as indicators of their resilience to disturbance and climate change (Kéfi *et al*., 2007, 2024; Veldhuis *et al*., 2022). For instance, in the GWW the patchy distribution of trees in old-growth stands can help prevent the spread of stand-replacing wildfires (Gosper *et al*., 2018; Jucker *et al*., 2023). But how these vegetation spatial patterns actually emerge as a result of variability in tree growth and survival rates across heterogeneous landscapes is not well established. In this regard our work provides a starting point for linking demography to vegetation spatial patterns through the integration of empirical data, remote sensing and models (Jucker, 2022).

Tracking the fate of individual trees through time also allowed us to partition aboveground biomass dynamics into gains and losses, and explicitly link these back to changes in vegetation 3D structure. Our results indicate that over the nine-year period of this study, the net balance between biomass gains from tree growth and losses due to mortality was positive across the landscape (Fig. 3). This strongly suggests that in the absence of large wildfires, these old-growth obligate-seeder woodlands can continue to operate as net carbon sinks even when several centuries old (Gosper *et al*., 2018). These positive biomass trends closely mirrored those in canopy cover and volume captured directly from the CHMs, both of which also increased across the landscape (Fig. 6e–f). Similar patterns of increasing woody cover have been reported in African savannas and woodlands over recent decades (Venter *et al*., 2018; Zhao *et al*., 2024), where increasing CO_2_ concentrations are believed to be shifting the balance in favour of trees overs C_4_ grasses.

Somewhat surprisingly, we found that net biomass change across the study site was positive even though a large disturbance event occurred sometime between 2012–2021 (Fig. 6). This event affected a strip approximately 0.6 km^2^ in size (2.4% of the landscape), within which a marked decrease in canopy cover, height and biomass were observed. Specifically, we estimate that as much as 1630 Mg of biomass were lost due high rates of tree mortality in the affected area, around 2.4% of the total woody biomass found across the entire site in 2012. As no large fires or human-related disturbances occurred in this period, this disturbance event was most likely the result of a tornado, which are common in the aftermath of tropical cyclones tracking inland from the north-western Australian coast. This provides clear evidence that other disturbances aside from stand-replacing fires can play an important role in shaping the structure and dynamics of these semi-arid woodlands (Yates *et al*., 1994).

### Tracking forest dynamics one tree at a time from above – future opportunities and challenges

From a methodological standpoint, an important contribution of our study is the development of an improved and more flexible implementation of the widely-used Dalponte & Coomes (2016) tree crown segmentation algorithm. By incorporating a two-stage crown delineation routine, we were able to substantially improve the retrieval of large trees that make up most of the aboveground biomass (Fig. 3a; Lutz *et al*., 2018). Stand-level estimates of tree density, basal area and aboveground biomass derived from our ALS-segmented trees closely matched independent observations from field data, suggesting our results are robust. Nonetheless, there are several ways in which our approach could be built on going forward. An obvious one would be to use complementary remote sensing approaches, such as canopy spectroscopy, to map the traits, functional groups or even species of the individually segmented crowns (Asner *et al*., 2015; Marconi *et al*., 2022; Beloiu *et al*., 2023). This would reveal how growth and mortality rates vary among dominant species, thus building a more complete picture of the dynamics of these ecosystems. From a demography perspective, the missing piece of the puzzle is quantifying tree recruitment. This would require more frequent ALS acquisitions to avoid underestimating recruitment rates due to high mortality of small trees. But even then, estimating recruitment from remote sensing will always remain a major challenge, especially in denser forests where seedlings recruit in the understory. For this, leveraging data from ground monitoring networks will continue to be essential.

More generally, it is only through the integration of novel remote sensing approaches with long-term field records that we will be able to develop more robust predictions of how forests are responding to rapid global change. There is great potential for harnessing these complementary data streams to build better individual-based models of forest dynamics (Lines *et al*., 2022). Traditionally these models have been parameterised and validated against field data, relying heavily on stem diameter measurements that only indirectly reflect whole-plant growth and competitive environment. Using repeat ALS data to map large numbers of individual trees allows us to capture population-level trends in growth and mortality in a much more comprehensive way, which is crucial for calibrating and validating forest models. This is particularly relevant for open canopy forests such as the GWW, where individual trees can be accurately detected in airborne and even satellite imagery (Brandt *et al*., 2020). In this regard, our study provides a blueprint for mobilising increasingly available high-resolution remote sensing data to the track the spatial and temporal dynamics of open canopy forests, so that we may better understand how they will respond in an increasingly warmer and more fire-prone future (Jucker *et al*., 2023).

## ACKNOWLEDGEMENTS

We acknowledge and pay our respects to the First Nations people in the Great Western Woodlands on whose land this work was conducted. The project was funded by a Western Australian State NRM grant to the Ngadju Conservation Aboriginal Corporation and CSIRO and supported by the Australian Government through the TERN Great Western Woodlands Supersite and the TERN Landscapes platform. Airborne LiDAR from 2021 was collected by Landgate (Western Australia’s land information authority) under the Capture WA program with additional support from the Western Australian Department of Biodiversity, Conservation and Attractions Goldfields Region office. TJ was supported by a UK NERC Independent Research Fellowship (grant: NE/S01537X/1).

## AUTHOR CONTRIBUTIONS

TJ and SMP conceived the idea for the project. KZ coordinated the 2021 ALS survey. RB led the processing and analysis of the data, with assistance from TJ and input from all co-authors. RB and TJ wrote the first draft of the manuscript, with all coauthors contributing substantially to revisions.

## DATA AVAILABILITY STATEMENT

The 2012 ALS data used in this study are openly available through the TERN data portal (https://portal.tern.org.au). All other data and code underpinning the results of this paper will be publicly archived on Zenodo following the review of this article and the corresponding DOI will be included upon publication. This includes: (i) 2012 and 2021 canopy height models (0.5-m resolution, GeoTIFF format), (ii) the digital terrain model and topographic wetness index rasters (5-m resolution, GeoTIFF format), and (iii) a data frame containing the tree-level demographic data, along with covariates included in the statistical models (CSV format).

## SUPPORTING INFORMATION

Additional supporting information may be found in the online version of this article.

**Fig. S1:** Study region

**Methods S1**: Masking out mulga scrub habitat

**Methods S2**: A new implementation of the *dalponte2016* segmentation algorithm

**Table S1:** Comparison of alternative tree segmentation algorithms

**Fig. S2:** Crown diameter to tree height allometries

**Fig. S3:** Crown detection accuracy of alternative tree segmentation algorithms

**Fig. S4:** Accuracy of alternative implementations of the *dalponte2016* algorithm

**Fig. S5:** Comparison of mean and maximum height change estimates

**Fig. S6:** Variation in ALS pulse density across the study area

**Fig. S7:** Shifts in height and crown growth allocation with tree size

## REFERENCES

Antin C, Le Bec J, Ayyappan N, Ramesh BR, Pélissier R. 2016. Allometric projections of time-related growth trajectories of two coexisting dipterocarp canopy species in India. Plant Ecology & Diversity 9: 603–614.

Asner GP, Martin RE, Anderson CB, Knapp DE. 2015. Quantifying forest canopy traits: Imaging spectroscopy versus field survey. Remote Sensing of Environment 158: 15–27.

Asner GP, Mascaro J. 2014. Mapping tropical forest carbon: Calibrating plot estimates to a simple LiDAR metric. Remote Sensing of Environment 140: 614–624.

Aubry-Kientz M, Dutrieux R, Ferraz A, Saatchi S, Hamraz H, Williams J, Coomes D, Piboule A, Vincent G. 2019. A comparative assessment of the performance of individual tree crowns delineation algorithms from ALS data in tropical forests. Remote Sensing 11: 1086.

Beese L, Dalponte M, Asner GP, Coomes DA, Jucker T. 2022. Using repeat airborne LiDAR to map the growth of individual oil palms in Malaysian Borneo during the 2015–16 El Niño. International Journal of Applied Earth Observation and Geoinformation 115: 103117.

Beloiu M, Heinzmann L, Rehush N, Gessler A, Griess VC. 2023. Individual tree-crown detection and species identification in heterogeneous forests using aerial RGB imagery and deep learning. Remote Sensing 15: 1463.

Boisvenue C, Running SW. 2006. Impacts of climate change on natural forest productivity – evidence since the middle of the 20th century. Global Change Biology 12: 862–882.

Brandt M, Gominski D, Reiner F, Kariryaa A, Guthula VB, Ciais P, Tong X, Zhang W, Govindarajulu D, Ortiz-Gonzalo D, et al. 2024. Severe decline in large farmland trees in India over the past decade. Nature Sustainability: 1–9.

Brandt M, Tucker CJ, Kariryaa A, Rasmussen K, Abel C, Small J, Chave J, Rasmussen LV, Hiernaux P, Diouf AA, et al. 2020. An unexpectedly large count of trees in the West African Sahara and Sahel. Nature 587: 78–82.

Canadell JG, Meyer CP (Mick), Cook GD, Dowdy A, Briggs PR, Knauer J, Pepler A, Haverd V. 2021. Multi-decadal increase of forest burned area in Australia is linked to climate change. Nature Communications 12: 6921.

Canham C, LePage P, Coates K. 2004. Canham CD, LePage PT, Coates KD. A neighborhood analysis of canopy tree competition: effects of shading versus crowding. Can J Forest Res 34: 778–787. Canadian Journal of Forest Research-Revue Canadienne De Recherche Forestiere 34: 778–787.

Cao Y, Ball JGC, Coomes DA, Steinmeier L, Knapp N, Wilkes P, Disney M, Calders K, Burt A, Lin Y, et al. 2023. Benchmarking airborne laser scanning tree segmentation algorithms in broadleaf forests shows high accuracy only for canopy trees. International Journal of Applied Earth Observation and Geoinformation 123: 103490.

Chambers JQ, Negron-Juarez RI, Marra DM, Di Vittorio A, Tews J, Roberts D, Ribeiro GHPM, Trumbore SE, Higuchi N. 2013. The steady-state mosaic of disturbance and succession across an old-growth Central Amazon forest landscape. Proceedings of the National Academy of Sciences 110: 3949–3954.

Choi DH, LaRue EA, Atkins JW, Foster JR, Matthes JH, Fahey RT, Thapa B, Fei S, Hardiman BS. 2023. Short-term effects of moderate severity disturbances on forest canopy structure. Journal of Ecology 111: 1866–1881.

Colgan MS, Asner GP, Levick SR, Martin RE, Chadwick OA. 2012. Topo-edaphic controls over woody plant biomass in South African savannas. Biogeosciences 9: 1809–1821.

Coomes DA, Allen RB. 2007. Mortality and tree-size distributions in natural mixed-age forests. Journal of Ecology 95: 27–40.

Coomes DA, Flores O, Holdaway R, Jucker T, Lines ER, Vanderwel MC. 2014. Wood production response to climate change will depend critically on forest composition and structure. Global Change Biology 20: 3632–3645.

Coomes DA, Holdaway RJ, Kobe RK, Lines ER, Allen RB. 2012. A general integrative framework for modelling woody biomass production and carbon sequestration rates in forests. Journal of Ecology 100: 42–64.

Cushman KC, Burley JT, Imbach B, Saatchi SS, Silva CE, Vargas O, Zgraggen C, Kellner JR. 2021. Impact of a tropical forest blowdown on aboveground carbon balance. Scientific Reports 11: 11279.

Cushman K c., Detto M, García M, Muller-Landau HC. 2022. Soils and topography control natural disturbance rates and thereby forest structure in a lowland tropical landscape. Ecology Letters 25: 1126–1138.

Dalagnol R, Wagner FH, Galvão LS, Streher AS, Phillips OL, Gloor E, Pugh TAM, Ometto JPHB, Aragão LEOC. 2021. Large-scale variations in the dynamics of Amazon forest canopy gaps from airborne lidar data and opportunities for tree mortality estimates. Scientific Reports 11: 1388.

Dalponte M, Coomes DA. 2016. Tree-centric mapping of forest carbon density from airborne laser scanning and hyperspectral data. Methods in Ecology and Evolution 7: 1236–1245.

Dalponte M, Jucker T, Liu S, Frizzera L, Gianelle D. 2019. Characterizing forest carbon dynamics using multi-temporal lidar data. Remote Sensing of Environment 224: 412–420.

Duncanson L, Dubayah R. 2018. Monitoring individual tree-based change with airborne lidar. Ecology and Evolution 8: 5079–5089.

Esquivel-Muelbert A, Phillips OL, Brienen RJW, Fauset S, Sullivan MJP, Baker TR, Chao K-J, Feldpausch TR, Gloor E, Higuchi N, et al. 2020. Tree mode of death and mortality risk factors across Amazon forests. Nature Communications 11: 5515.

Eysn L, Hollaus M, Lindberg E, Berger F, Monnet J-M, Dalponte M, Kobal M, Pellegrini M, Lingua E, Mongus D, et al. 2015. A benchmark of lidar-based single tree detection methods using heterogeneous forest data from the Alpine Space. Forests 6: 1721–1747.

Ferraz A, Saatchi S, Mallet C, Meyer V. 2016. Lidar detection of individual tree size in tropical forests. Remote Sensing of Environment 183: 318–333.

Fischer FJ, Labrière N, Vincent G, Hérault B, Alonso A, Memiaghe H, Bissiengou P, Kenfack D, Saatchi S, Chave J. 2020. A simulation method to infer tree allometry and forest structure from airborne laser scanning and forest inventories. Remote Sensing of Environment 251: 112056.

Fischer FJ, Maréchaux I, Chave J. 2019. Improving plant allometry by fusing forest models and remote sensing. New Phytologist 223: 1159–1165.

Fisher RA, Koven CD, Anderegg WRL, Christoffersen BO, Dietze MC, Farrior CE, Holm JA, Hurtt GC, Knox RG, Lawrence PJ, et al. 2018. Vegetation demographics in Earth System Models: A review of progress and priorities. Global Change Biology 24: 35–54.

Gosper CR, Yates CJ, Cook GD, Harvey JM, Liedloff AC, McCaw WL, Thiele KR, Prober SM. 2018. A conceptual model of vegetation dynamics for the unique obligate-seeder eucalypt woodlands of south-western Australia. Austral Ecology 43: 681–695.

Hammond WM, Williams AP, Abatzoglou JT, Adams HD, Klein T, López R, Sáenz-Romero C, Hartmann H, Breshears DD, Allen CD. 2022. Global field observations of tree die-off reveal hotter-drought fingerprint for Earth’s forests. Nature Communications 13: 1761.

Hartmann H, Bastos A, Das AJ, Esquivel-Muelbert A, Hammond WM, Martínez-Vilalta J, McDowell NG, Powers JS, Pugh TAM, Ruthrof KX, et al. 2022. Climate change risks to global forest health: emergence of unexpected events of elevated tree mortality worldwide. Annual Review of Plant Biology 73: 673–702.

Hijmans RJ, Bivand R, Dyba K, Pebesma E, Sumner MD. 2023. terra: Spatial Data Analysis. v.1.7-78. URL https://CRAN.R-project.org/package=terra.

Holcomb A, Mathis SV, Coomes DA, Keshav S. 2023. Computational tools for assessing forest recovery with GEDI shots and forest change maps. Science of Remote Sensing 8: 100106.

Hubau W, Lewis SL, Phillips OL, Affum-Baffoe K, Beeckman H, Cuní-Sanchez A, Daniels AK, Ewango CEN, Fauset S, Mukinzi JM, et al. 2020. Asynchronous carbon sink saturation in African and Amazonian tropical forests. Nature 579: 80–87.

Johnson MO, Galbraith D, Gloor M, De Deurwaerder H, Guimberteau M, Rammig A, Thonicke K, Verbeeck H, von Randow C, Monteagudo A, et al. 2016. Variation in stem mortality rates determines patterns of above-ground biomass in Amazonian forests: implications for dynamic global vegetation models. Global Change Biology 22: 3996–4013.

Jucker T. 2022. Deciphering the fingerprint of disturbance on the three-dimensional structure of the world’s forests. New Phytologist 233: 612–617.

Jucker T, Bongalov B, Burslem DFRP, Nilus R, Dalponte M, Lewis SL, Phillips OL, Qie L, Coomes DA. 2018. Topography shapes the structure, composition and function of tropical forest landscapes. Ecology Letters 21: 989–1000.

Jucker T, Bouriaud O, Coomes DA. 2015. Crown plasticity enables trees to optimize canopy packing in mixed-species forests. Functional Ecology 29: 1078–1086.

Jucker T, Caspersen J, Chave J, Antin C, Barbier N, Bongers F, Dalponte M, van Ewijk KY, Forrester DI, Haeni M, et al. 2017. Allometric equations for integrating remote sensing imagery into forest monitoring programmes. Global Change Biology 23: 177–190.

Jucker T, Fischer FJ, Chave J, Coomes DA, Caspersen J, Ali A, Loubota Panzou GJ, Feldpausch TR, Falster D, Usoltsev VA, et al. 2022. Tallo: A global tree allometry and crown architecture database. Global Change Biology 28: 5254–5268.

Jucker T, Gosper CR, Wiehl G, Yeoh PB, Raisbeck-Brown N, Fischer FJ, Graham J, Langley H, Newchurch W, O’Donnell AJ, et al. 2023. Using multi-platform LiDAR to guide the conservation of the world’s largest temperate woodland. Remote Sensing of Environment 296: 113745.

Kéfi S, Génin A, Garcia-Mayor A, Guirado E, Cabral JS, Berdugo M, Guerber J, Solé R, Maestre FT. 2024. Self-organization as a mechanism of resilience in dryland ecosystems. Proceedings of the National Academy of Sciences 121: e2305153121.

Kéfi S, Rietkerk M, Alados CL, Pueyo Y, Papanastasis VP, ElAich A, de Ruiter PC. 2007. Spatial vegetation patterns and imminent desertification in Mediterranean arid ecosystems. Nature 449: 213–217.

Khosravipour A, Skidmore A, Isenburg M, Hussin Y. 2014. Generating pit-free canopy height models from airborne lidar. Photogrammetric Engineering & Remote Sensing 80: 863– 872.

Kopecký M, Macek M, Wild J. 2021. Topographic Wetness Index calculation guidelines based on measured soil moisture and plant species composition. Science of The Total Environment 757: 143785.

Kunstler G, Guyennon A, Ratcliffe S, Rüger N, Ruiz-Benito P, Childs DZ, Dahlgren J, Lehtonen A, Thuiller W, Wirth C, et al. 2021. Demographic performance of European tree species at their hot and cold climatic edges. Journal of Ecology 109: 1041–1054.

Laurans M, Munoz F, Charles-Dominique T, Heuret P, Fortunel C, Isnard S, Sabatier S- A, Caraglio Y, Violle C. 2024. Why incorporate plant architecture into trait-based ecology? Trends in Ecology & Evolution 0.

Lines ER, Coomes DA, Purves DW. 2010. Influences of Forest Structure, Climate and Species Composition on Tree Mortality across the Eastern US. PLOS ONE 5: e13212.

Lines ER, Fischer FJ, Owen HJF, Jucker T. 2022. The shape of trees: Reimagining forest ecology in three dimensions with remote sensing. Journal of Ecology 110: 1730–1745.

Lutz JA, Furniss TJ, Johnson DJ, Davies SJ, Allen D, Alonso A, Anderson-Teixeira KJ, Andrade A, Baltzer J, Becker KML, et al. 2018. Global importance of large-diameter trees. Global Ecology and Biogeography 27: 849–864.

Ma Q, Su Y, Niu C, Ma Q, Hu T, Luo X, Tai X, Qiu T, Zhang Y, Bales RC, et al. 2023. Tree mortality during long-term droughts is lower in structurally complex forest stands. Nature Communications 14: 7467.

Marconi S, Weinstein BG, Zou S, Bohlman SA, Zare A, Singh A, Stewart D, Harmon I, Steinkraus A, White EP. 2022. Continental-scale hyperspectral tree species classification in the United States National Ecological Observatory Network. Remote Sensing of Environment 282: 113264.

Marziliano PA, Tognetti R, Lombardi F. 2019. Is tree age or tree size reducing height increment in Abies alba Mill. at its southernmost distribution limit? Annals of Forest Science 76: 1–12.

McDowell NG, Allen CD, Anderson-Teixeira K, Aukema BH, Bond-Lamberty B, Chini L, Clark JS, Dietze M, Grossiord C, Hanbury-Brown A, et al. 2020. Pervasive shifts in forest dynamics in a changing world. *Science (New York*, N.Y.) 368: eaaz9463.

Metcalfe P, Beven K, Freer J. 2015. Dynamic TOPMODEL: A new implementation in R and its sensitivity to time and space steps. Environmental Modelling & Software 72: 155–172.

Moncrieff GR, Chamaillé-Jammes S, Higgins SI, O’Hara RB, Bond WJ. 2011. Tree allometries reflect a lifetime of herbivory in an African savanna. Ecology 92: 2310–2315.

Muscarella R, Kolyaie S, Morton DC, Zimmerman JK, Uriarte M. 2020. Effects of topography on tropical forest structure depend on climate context. Journal of Ecology 108: 145–159.

Needham JF, Arellano G, Davies SJ, Fisher RA, Hammer V, Knox RG, Mitre D, Muller-Landau HC, Zuleta D, Koven CD. 2022a. Tree crown damage and its effects on forest carbon cycling in a tropical forest. Global Change Biology 28: 5560–5574.

Needham JF, Johnson DJ, Anderson-Teixeira KJ, Bourg N, Bunyavejchewin S, Butt N, Cao M, Cárdenas D, Chang-Yang C-H, Chen Y-Y, et al. 2022b. Demographic composition, not demographic diversity, predicts biomass and turnover across temperate and tropical forests. Global Change Biology 28: 2895–2909.

Nunes MH, Jucker T, Riutta T, Svátek M, Kvasnica J, Rejžek M, Matula R, Majalap N, Ewers RM, Swinfield T, et al. 2021. Recovery of logged forest fragments in a human-modified tropical landscape during the 2015-16 El Niño. Nature Communications 12: 1526.

Olson ME, Soriano D, Rosell JA, Anfodillo T, Donoghue MJ, Edwards EJ, León-Gómez C, Dawson T, Camarero Martínez JJ, Castorena M, et al. 2018. Plant height and hydraulic vulnerability to drought and cold. Proceedings of the National Academy of Sciences 115: 7551– 7556.

Paul KI, Roxburgh SH, Chave J, England JR, Zerihun A, Specht A, Lewis T, Bennett LT, Baker TG, Adams MA, et al. 2016. Testing the generality of above-ground biomass allometry across plant functional types at the continent scale. Global Change Biology 22: 2106–2124.

Pebesma E. 2018. Simple Features for R: Standardized Support for Spatial Vector Data. The R Journal 10: 439–446.

Piponiot C, Anderson-Teixeira KJ, Davies SJ, Allen D, Bourg NA, Burslem DFRP, Cárdenas D, Chang-Yang C-H, Chuyong G, Cordell S, et al. 2022. Distribution of biomass dynamics in relation to tree size in forests across the world. New Phytologist 234: 1664–1677.

Prober SM, Wiehl G, Gosper CR, Schultz L, Langley H, Macfarlane C. 2023. The Great Western Woodlands TERN SuperSite: ecosystem monitoring infrastructure and key science learnings. Journal of Ecology and Environment 47.

Pugh TAM, Rademacher T, Shafer SL, Steinkamp J, Barichivich J, Beckage B, Haverd V, Harper A, Heinke J, Nishina K, et al. 2020. Understanding the uncertainty in global forest carbon turnover. Biogeosciences 17: 3961–3989.

R Core Team. 2024. R: A language and environment for statistical computing. Vienna, Austria: R Foundation for Statistical Computing.

Rodriguez-Iturbe I, Chen Z, Staver AC, Levin SA. 2019. Tree clusters in savannas result from islands of soil moisture. Proceedings of the National Academy of Sciences 116: 6679– 6683.

Roussel J-R, Auty D, Coops NC, Tompalski P, Goodbody TRH, Meador AS, Bourdon J- F, de Boissieu F, Achim A. 2020. lidR: An R package for analysis of Airborne Laser Scanning (ALS) data. Remote Sensing of Environment 251: 112061.

Roussel J-R, Caspersen J, Béland M, Thomas S, Achim A. 2017. Removing bias from LiDAR-based estimates of canopy height: Accounting for the effects of pulse density and footprint size. Remote Sensing of Environment 198: 1–16.

Ruiz-Benito P, Lines ER, Gómez-Aparicio L, Zavala MA, Coomes DA. 2013. Patterns and drivers of tree mortality in Iberian forests: Climatic effects are modified by competition. PLOS ONE 8: e56843.

Senf C, Pflugmacher D, Zhiqiang Y, Sebald J, Knorn J, Neumann M, Hostert P, Seidl R. 2018. Canopy mortality has doubled in Europe’s temperate forests over the last three decades. Nature Communications 9: 4978.

Stephenson NL, Das AJ, Condit R, Russo SE, Baker PJ, Beckman NG, Coomes DA, Lines ER, Morris WK, Rüger N, et al. 2014. Rate of tree carbon accumulation increases continuously with tree size. Nature 507: 90–93.

Stovall AEL, Shugart H, Yang X. 2019. Tree height explains mortality risk during an intense drought. Nature Communications 10: 4385.

Swetnam TL, Brooks PD, Barnard HR, Harpold AA, Gallo EL. 2017. Topographically driven differences in energy and water constrain climatic control on forest carbon sequestration. Ecosphere 8: e01797.

Taubert F, Jahn MW, Dobner H-J, Wiegand T, Huth A. 2015. The structure of tropical forests and sphere packings. Proceedings of the National Academy of Sciences 112: 15125– 15129.

Turner MG, Seidl R. 2023. Novel Disturbance Regimes and Ecological Responses. Annual Review of Ecology, Evolution, and Systematics 54: 63–83.

Veldhuis MP, Martinez-Garcia R, Deblauwe V, Dakos V. 2022. Remotely-sensed slowing down in spatially patterned dryland ecosystems. Ecography 2022: e06139.

Venter ZS, Cramer MD, Hawkins H-J. 2018. Drivers of woody plant encroachment over Africa. Nature Communications 9: 2272.

Wang Y, Hyyppä J, Liang X, Kaartinen H, Yu X, Lindberg E, Holmgren J, Qin Y, Mallet C, Ferraz A, et al. 2016. International benchmarking of the individual tree detection methods for modelling 3-D canopy structure for silviculture and forest ecology using Airborne Laser Scanning. IEEE Transactions on Geoscience and Remote Sensing 54: 5011–5027.

Wedeux B, Dalponte M, Schlund M, Hagen S, Cochrane M, Graham L, Usup A, Thomas A, Coomes D. 2020. Dynamics of a human-modified tropical peat swamp forest revealed by repeat lidar surveys. Global Change Biology 26: 3947–3964.

Yates CJ, Hobbs RJ, Bell RW. 1994. Landscape-scale disturbances and regeneration in semi-arid woodlands of southwestern Australia. Pacific Conservation Biology 1: 214–221.

Yu K, Chen HYH, Gessler A, Pugh TAM, Searle EB, Allen RB, Pretzsch H, Ciais P, Phillips OL, Brienen RJW, et al. 2024. Forest demography and biomass accumulation rates are associated with transient mean tree size vs. density scaling relations. PNAS Nexus 3: pgae008.

Yu Y, Saatchi S, Domke GM, Walters B, Woodall C, Ganguly S, Li S, Kalia S, Park T, Nemani R, et al. 2022. Making the US national forest inventory spatially contiguous and temporally consistent. Environmental Research Letters 17: 065002.

Zhao Z, Ciais P, Wigneron J-P, Santoro M, Brandt M, Kleinschroth F, Lewis SL, Chave J, Fensholt R, Laporte N, et al. 2024. Central African biomass carbon losses and gains during 2010–2019. One Earth 7: 506–519.

Zuidema PA, van der Sleen P. 2022. Seeing the forest through the trees: how tree-level measurements can help understand forest dynamics. New Phytologist 234: 1544–1546.

Zuleta D, Arellano G, Muller-Landau HC, McMahon SM, Aguilar S, Bunyavejchewin S, Cárdenas D, Chang-Yang C-H, Duque A, Mitre D, et al. 2022. Individual tree damage dominates mortality risk factors across six tropical forests. New Phytologist 233: 705–721.

