## Supporting information for "Tracking tree demography and forest dynamics at scale using remote sensing"

### Fig. S1: Study region

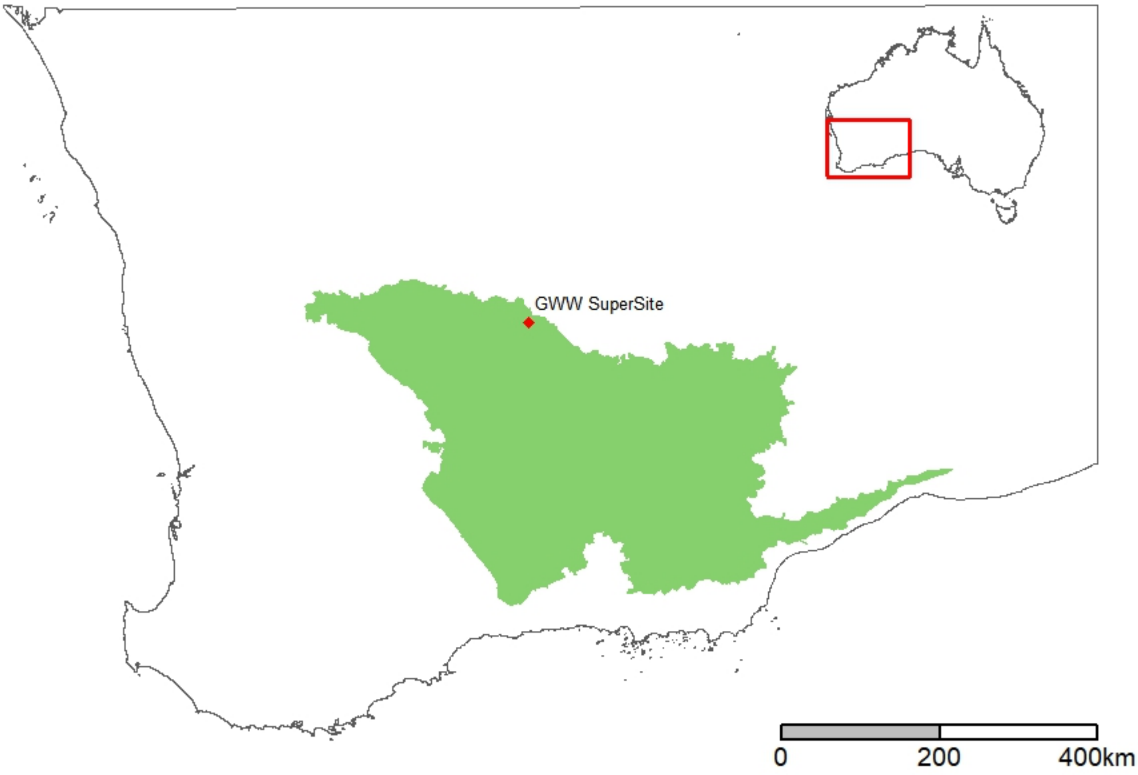

**Fig. S1**: Location of the TERN SuperSite within the Great Western Woodlands (green shaded area) and in relation to the rest of Australia (inset map).

### Methods S1: Masking out mulga scrub habitat

Our study focused on the dynamics of obligate-seeded eucalypt woodlands, which dominate much of the TERN SuperSite. However, our study site also included small, scattered patches of *Acacia aneura* (commonly known as mulga), which are shorter and denser than the surrounding obligate-seeded woodland. To avoid these impacting our results, we developed a semi-automated approach to mask out these areas of mulga. Specifically, mulga vegetation are considerably denser that the surrounding habitat, despite being shorter. We therefore used reference parameters of canopy cover measured from the CHM in an area known to be dominated by mulga to set a threshold which we applied across the entire landscape (defined as >60% ground cover at 2m aboveground over a contiguous area of at least 40×40 m). We then used the RGB imagery acquired in 2021 to manually screen areas of landscape that met this threshold, excluding those that were structurally and spectrally consistent with mulga habitat. In total, this led us to masking out a cumulative area of 9 ha (0.4% of the total) from all subsequent analyses.

### Methods S2: A new implementation of the *dalponte2016* segmentation algorithm

The *dalponte2016* algorithm works by first applying a local maximum filter (LMF) to locate the tops of individual trees in the CHM and then uses these as starting points around which to delineate the borders of their crowns following a set of rules. A key feature of *dalponte2016* is that the size of the window within which the LMF searches for treetops can be allowed to vary depending on the height of the canopy (Coomes *et al.*, 2017), reflecting underlying allometric constraints between crown diameter (*CD*, in m) and tree height (*H*, in m). For the default implementation of the *dalponte2016* algorithm, we used data from the 797 manually delineated tree crowns to model *CD* as a power-law function of *H* by fitting a linear model to log–log transformed data (Jucker *et al.*, 2022). This yielded the following power-law function: $CD = 0.957 \times H^{0.988}$ (Eqn. 1). However, as previous studies have shown, this approach is prone to over-segmenting the crowns of large, canopy dominant trees (Cao *et al.*, 2023). Given that these large trees play a disproportionately important role in shaping forest biomass dynamics, it is crucial to segment their crowns as well as possible. One way to do this is to use quantile regression to model the upper percentiles of the *CD*–*H* data (e.g., 95$^{th}$ percentile), thus effectively widening the search window within which the LMF searches for treetops and reducing the likelihood of over-segmentation (Cao *et al.*, 2023). However, this typically then comes at the expense of missing larger numbers of smaller crowns (i.e,, high rates of omission). To get the best of both approaches, we therefore developed a two-stage segmentation routine. The first step involved using quantile regression to set the size of the search window, for which we tested models fit to the upper 70–95% percentiles of the data in 5% increments. We then overlapped these crowns with those identified using Eqn. 1 and retained any that had been missed when using quantile regression to set the search window. Based on this we determined that for our study the optimal combination was achieved when using an initial search window defined by the 80$^{th}$ percentile of the data ($CD = 2.040 \times H^{0.764}$; Table S1 and Fig. S2).

### Table S1: Comparison of alternative tree segmentation algorithms

**Table S1**: We compared alternative crown segmentation algorithms implemented in the *lidR* package against a set of 797 manually delineated tree crowns. The performance of each algorithm was assessed on the basis of the following criteria: (i) total number of segmented crowns, (ii) mean crown area of correctly matched crowns, (iii) proportion of correctly segmented crowns, (iv) proportion of over-segmented crowns, (v) proportion of omitted crowns, (vi) intersection over union (IoU) of matched crows, (vii) precision, (viii) recall, and (ix) $F_1$ score. For *dalponte2016*, we compared the performance of the algorithm implemented using the default search window – which characteries the expected relationship between crown diameter and tree height using a linear regression model (lm) – to a customised version in which we combined the lm search window with a second search window defined using quantile regression (qr) fit to various different percentiles of the data (70–95$^{th}$ percentile). Fig. S2 below illustrates the differences between the lm and qr approach. Figs S3-4 provide a visual comparison of the performance of the various algorithms described below.

| **Algorithm** | **Number of crowns** | **Mean crown area ($m^2$)** | **Correctly segmented (%)** | **Over-segmented (%)** | **Omitted (%)** | **IoU (%)** | **Precision** | **Recall** | **$F_1$ score** |
|---|---|---|---|---|---|---|---|---|---|
| Manual reference dataset | 797 | 132 | | | | | | | |
| *dalponte2016* (lm) | 932 | 128 | 78 | 13 | 9 | 75 | 0.89 | 0.96 | 0.92 |
| *watershed* | 789 | 157 | 80 | 5 | 15 | 68 | 0.73 | 0.97 | 0.84 |
| *li2012* | 2393 | 173 | 40 | 50 | 10 | 32 | 0.76 | 0.98 | 0.86 |
| *silva2016* | 984 | 84 | 81 | 12 | 7 | 62 | 0.07 | 1.5 | 0.13 |
| *dalponte2016* (lm + $qr_{95}$) | 774 | 134 | 81 | 3 | 16 | 72 | 0.82 | 0.95 | 0.88 |
| *dalponte2016* (lm + $qr_{90}$) | 796 | 132 | 81 | 4 | 15 | 73 | 0.83 | 0.95 | 0.89 |
| *dalponte2016* (lm + $qr_{85}$) | 806 | 131 | 81 | 5 | 14 | 73 | 0.84 | 0.95 | 0.90 |
| *dalponte2016* (lm + $qr_{80}$) | 825 | 130 | 82 | 5 | 13 | 75 | 0.86 | 0.95 | 0.91 |
| *dalponte2016* (lm + $qr_{75}$) | 832 | 130 | 83 | 5 | 12 | 75 | 0.86 | 0.95 | 0.91 |
| *dalponte2016* (lm + $qr_{70}$) | 841 | 129 | 82 | 6 | 12 | 75 | 0.86 | 0.95 | 0.91 |

**Fig. S2: Crown diameter to tree height allometries**

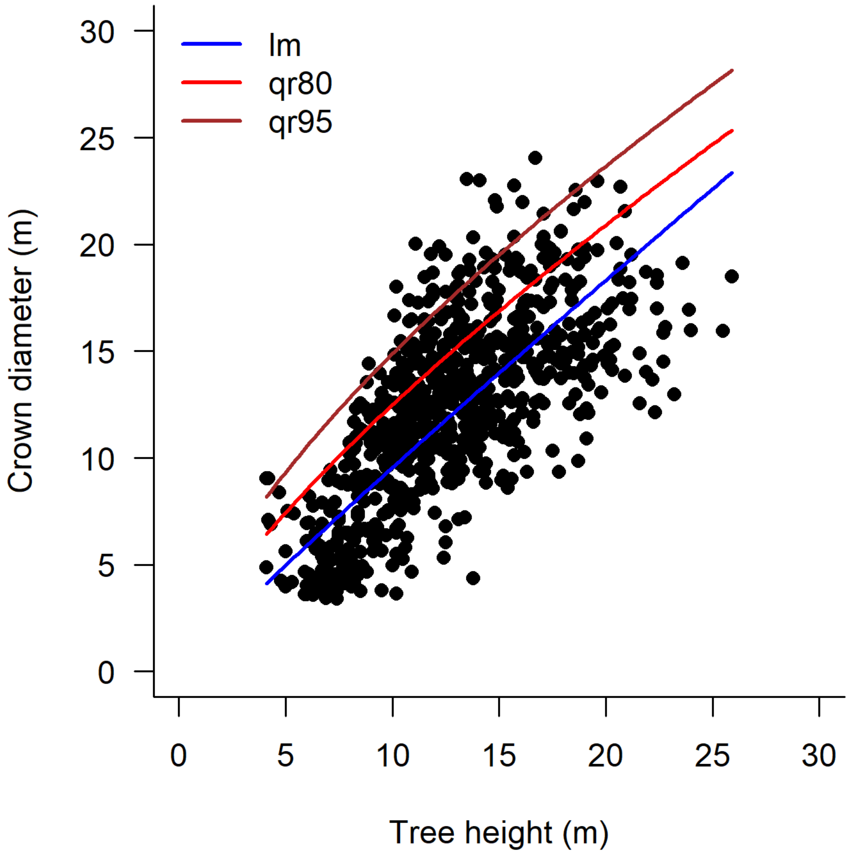

**Fig. S2**: Relationship between crown diameter and tree height derived from the 797 manually delineated tree crowns. Curves show the fit of alternative allometric models relating the two axes of crown size, including a linear regression model (lm, blue curve) and a quantile regression model fit to the 80th (qr80, red curve) and 95th (qr95, brown curve) percentile of the data. All three models were fit to log-log transformed data.

### Fig. S3: Crown detection accuracy of alternative tree segmentation algorithms

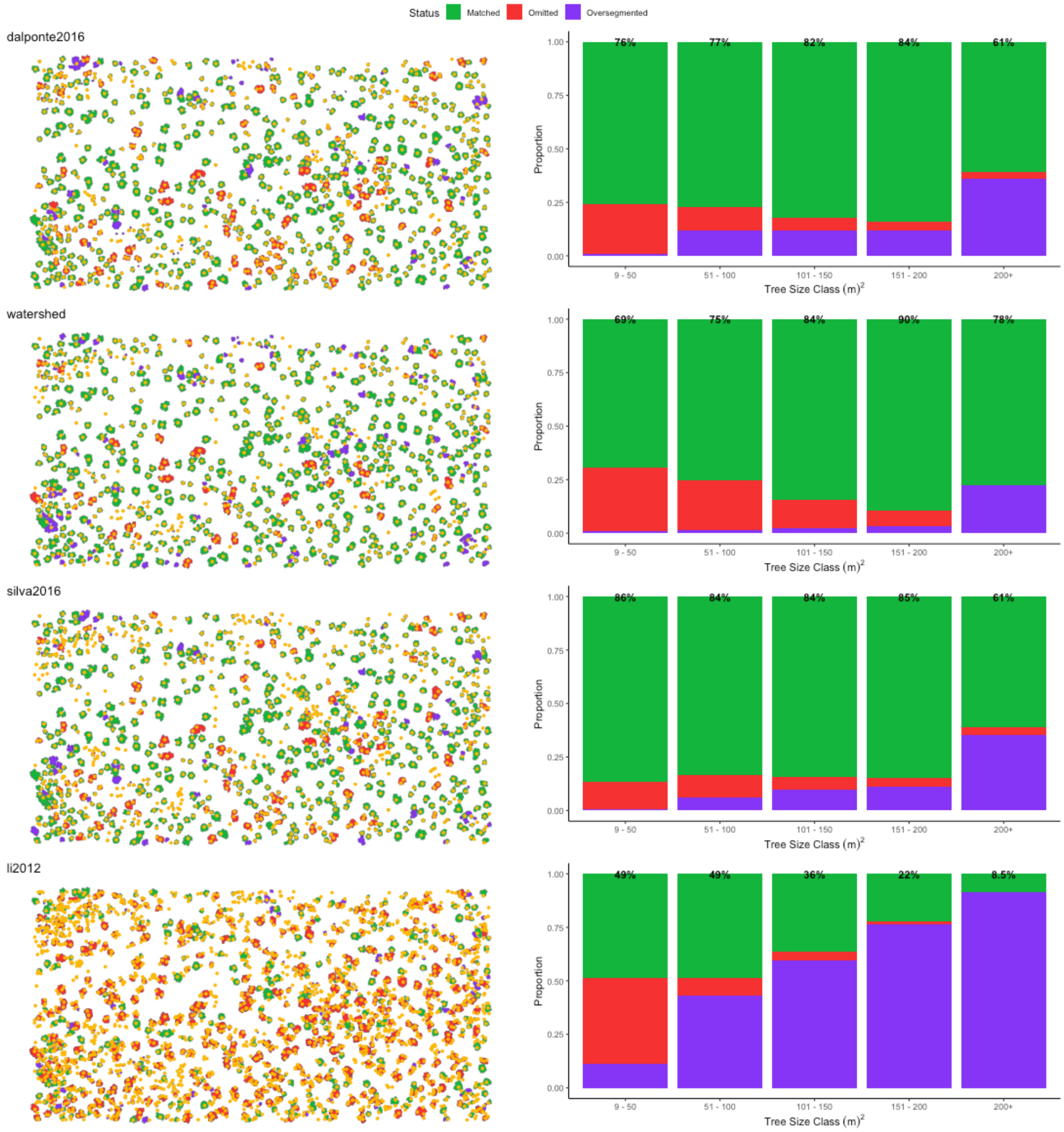

**Fig. S3**: Comparison of the accuracy of the four crown delineation algorithms tested in this study (from top to bottom: *dalponte2016*, *watershed*, *silva2016* and *li2012*; see Table S1 for details). For each algorithm, the panel on the left shows a map of the 797 manually delineated tree crowns used to assess the accuracy of the delineation, overlapped onto which are the centroids of the crowns identified by the respective algorithm (yellow dots). Those in green represent correctly matched crowns, those in red were omitted by the algorithm, and those in purple were over-segmented. The barplots on the right summarise the results of the segmentation algorithms for trees of different size classes (defined here on the basis of their crown area). The values reported at the top of each bar correspond to the percentage of trees correctly segmented in each size class.

### Fig. S4: Accuracy of alternative implementations of the *dalponte2016* algorithm

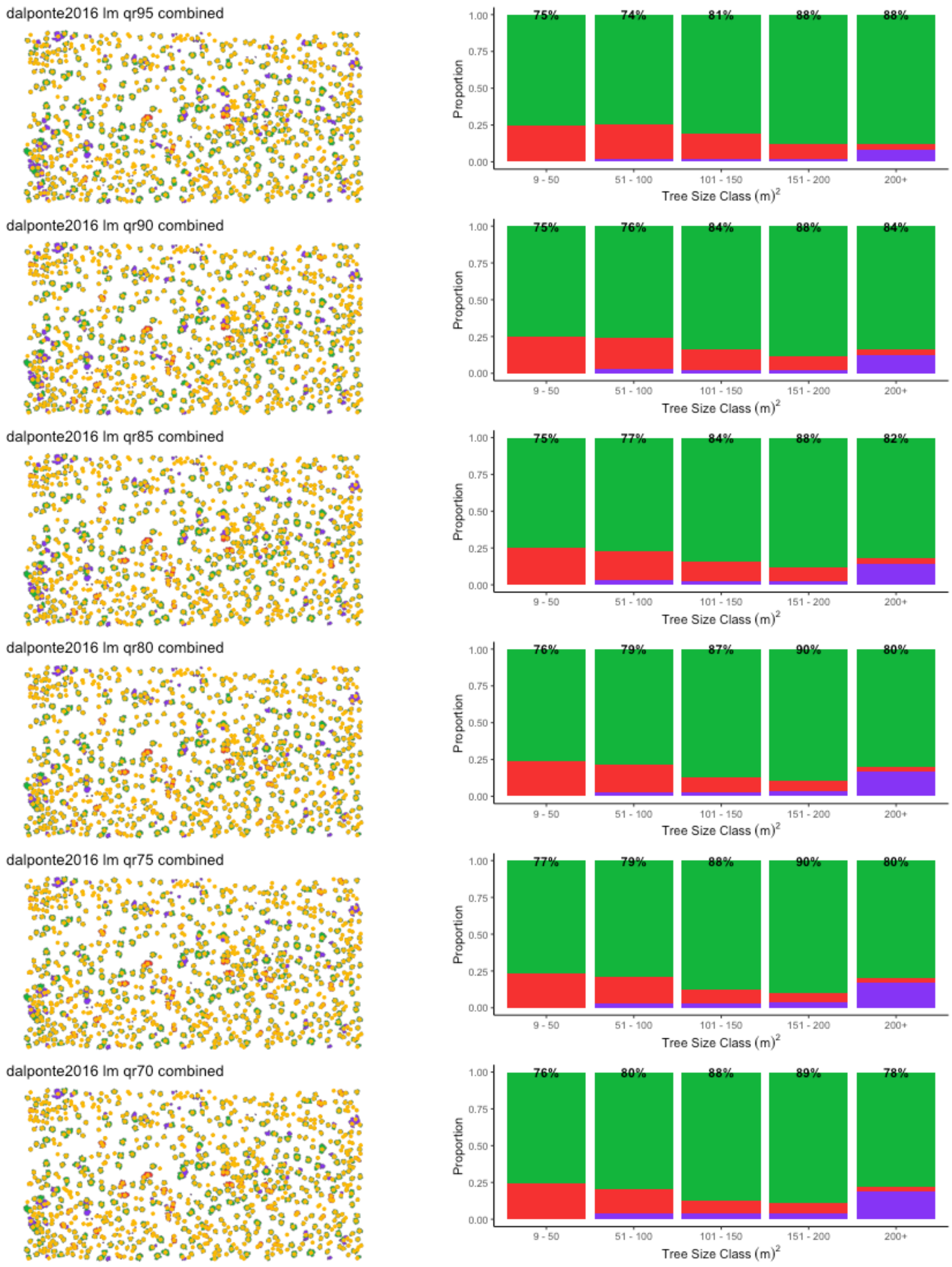

**Fig. S4**: Comparison of the accuracy of different iterations of the *dalponte2016* segmentation algorithm in which we combined a search window defined the parameters of a linear regression model (lm) relating crown diameter to tree height (blue curve in Fig. S2) with a second search window defined using quantile regression (qr) fit to various different percentiles of the data (70–95th percentile). For each combination, the panel on the left shows a map of the 797 manually delineated tree crowns used to assess the accuracy of the delineation, overlapped onto which are the centroids of the crowns identified by the respective algorithm (yellow dots). Those in green represent correctly matched crowns, those in red were omitted by the algorithm, and those in purple were over-segmented. The barplots on the right summarise the results of the segmentation algorithms for trees of different size classes (defined here on the basis of their crown area). The values reported at the top of each bar correspond to the percentage of trees correctly segmented in each size class.

**Fig. S5: Comparison of mean and maximum height change estimates**

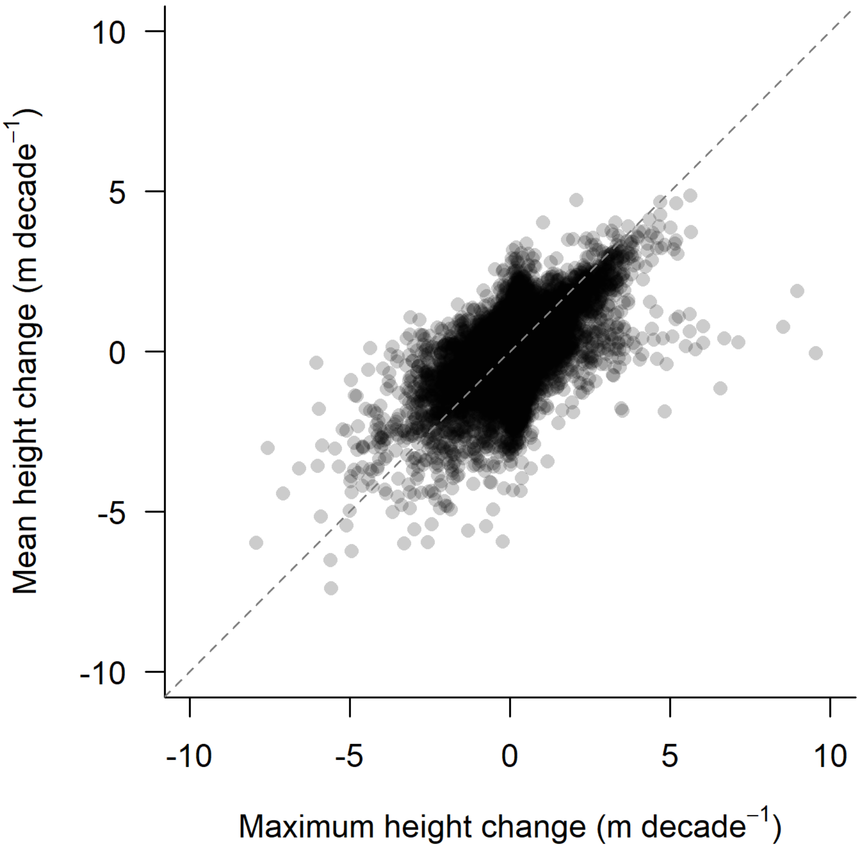

**Fig. S5**: Comparison of individual tree height change estimates (n = 39,286 trees) based on mean crown height (i.e., mean value of CHM pixels falling inside the crown polygon) and maximum crown height (i.e., maximum value of CHM pixels falling inside the crown polygon). The two estimates of tree height change were strongly correlated (Pearson's correlation coefficient = 0.53; $P < 0.0001$) and were similar in terms of absolute values (dashed grey line indicates a 1:1 relationship).

### Fig. S6: Variation in ALS pulse density across the study area

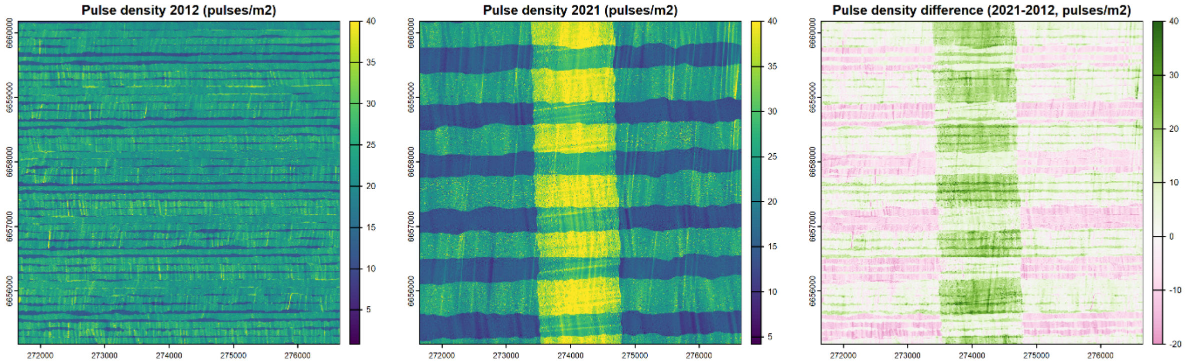

**Fig. S6**: Maps showing variation in pulse density in (**a**) 2012 and (**b**) 2021 across the TERN SuperSite, as well as (**c**) the differences between the two ALS surveys. The resolution of the maps is 5×5 m.

**Fig. S7: Shifts in height and crown growth allocation with tree size**

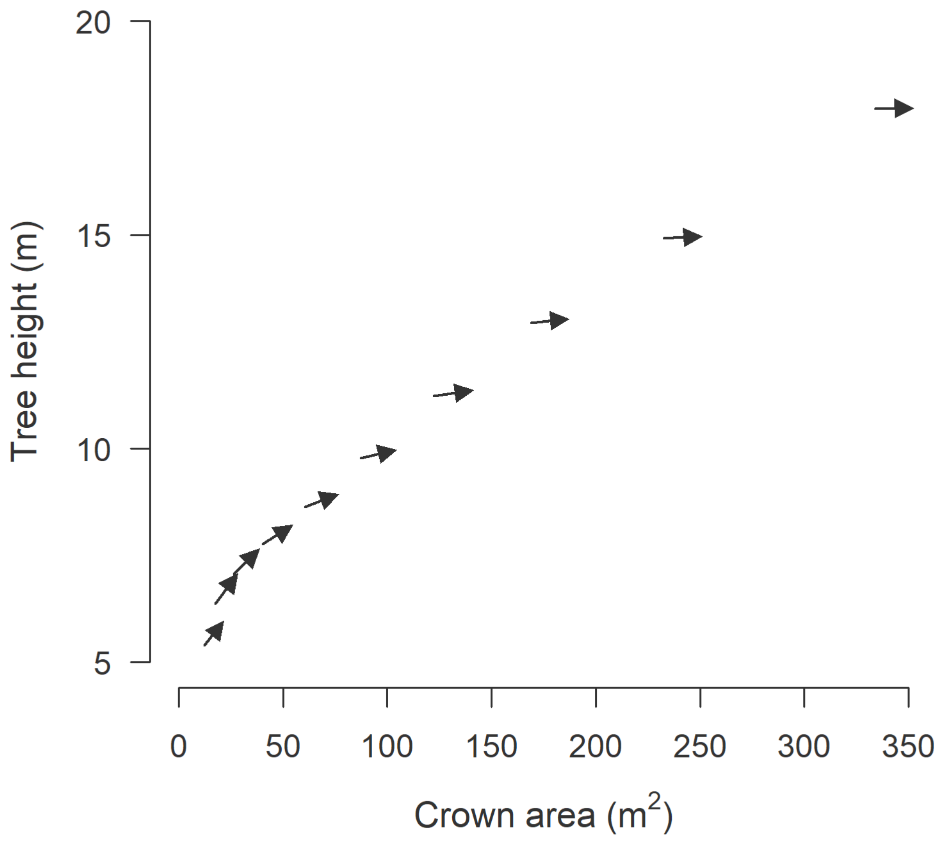

**Fig. S7**: Changes in allocation to height and crown area growth with tree size. Each arrow illustrates the growth trajectory of trees within a size class, with the beginning of the arrow corresponding to the mean height and crown area of trees in 2012 and the end of the arrow indicating the size reached by the time of the second survey in 2021. Arrows pointing upwards at a 45° angle indicate trees growing both in height and crown area, while ones pointing horizontally to the right denote trees that expanded their crowns but remained unchanged in terms of height. Trees were grouped into 10 equal width size class bins, where size was defined as the as the product of tree height and crown diameter. Each size class in the analysis is represented by over 1750 trees.

### References

Cao Y, Ball JGC, Coomes DA, Steinmeier L, Knapp N, Wilkes P, Disney M, Calders K, Burt A, Lin Y, *et al.* 2023. Benchmarking airborne laser scanning tree segmentation algorithms in broadleaf forests shows high accuracy only for canopy trees. *International Journal of Applied Earth Observation and Geoinformation* 123: 103490.

Coomes DA, Dalponte M, Jucker T, Asner GP, Banin LF, Burslem DFRP, Lewis SL, Nilus R, Phillips OL, Phua M-H, *et al.* 2017. Area-based vs tree-centric approaches to mapping forest carbon in Southeast Asian forests from airborne laser scanning data. *Remote Sensing of Environment* 194: 77–88.

Jucker T, Fischer FJ, Chave J, Coomes DA, Caspersen J, Ali A, Loubota Panzou GJ, Feldpausch TR, Falster D, Usoltsev VA, *et al.* 2022. Tallo: A global tree allometry and crown architecture database. *Global Change Biology* 28: 5254–5268.
